# The JEDI marker as a universal measure of planetary biodiversity

**DOI:** 10.1101/2025.08.11.669668

**Authors:** Taylor Priest, Nicolas Henry, Thomas Weber, Laurine Planat, Coralie Rousseau, Simon M. Dittami, Yi-Chun Yeh, David M. Needham, Hans-Joachim Ruscheweyh, Fabienne Rigaut-Jalabert, Nathalie Simon, Sarah Romac, Florence Le Gall, Thomas Beavis, Kevin Moog, Alice Moussy, Corinne Da Silva, Caroline Belser, ECOMAP team, TREC expedition team, Mobile Laboratory team, Jan Korbel, Raffaele Siano, Julie Poulain, Patrick Wincker, Paola Bertucci, Peer Bork, Jed A. Fuhrman, Flora Vincent, Shinichi Sunagawa, Colomban de Vargas

**Affiliations:** Department of Biology, Institute of Microbiology and Swiss Institute of Bioinformatics, ETH Zurich, Zurich, Switzerland; Research Federation for the study of Global Ocean Systems Ecology and Evolution, FR2022 Tara GOSEE, F-75016 Paris, France; CNRS, Sorbonne Université, FR2424, ABiMS, Station Biologique de Roscoff, Roscoff, France; Data Science Centre, European Molecular Biology Laboratory, Heidelberg, Germany; Developmental Biology Unit, European Molecular Biology Laboratory, 69117 Heidelberg, Germany; Integrative Biology of Marine Models (LBI2M, UMR 8227), Station Biologique de Roscoff, Sorbonne Université, CNRS, Roscoff, France; Institute of Oceanography, National Taiwan University, Taipei, Taiwan; GEOMAR Helmholtz Centre for Ocean Research Kiel, Kiel, Germany; Faculty of Mathematics and Natural Sciences, Kiel University, Kiel, Germany; Sorbonne Université, CNRS, Adaptation et Diversité en Milieu Marin, AD2M, F-29680 Roscoff, France; Genoscope, Institut de Biologie François Jacob, Commissariat à l’énergie atomique (CEA), CNRS, Université de Paris-Saclay Evry 91057 France; IFREMER, DYNECO, BP70, Plouzané, France; European Molecular Biology Laboratory, EMBL Barcelona, C/ Dr. Aiguader, 88, PRBB Building, 08003 Barcelona, Spain; European Molecular Biology Laboratory, Meyerhofstraße 1, 69117 Heidelberg, Germany; Department of Biological Sciences, University of Southern California, Los Angeles, USA

**Author notes:** Authors have contributed equally (Taylor Priest, Nicolas Henry).

## Abstract

Despite its critical importance in the formation and maintenance of ecosystems and homeostasis on Earth, biodiversity remains a complex and non-unified concept. Consequently, standards for measuring global biodiversity are lacking, hindering our capacity to document Earth’s biota and track its change. Here, we propose the ‘Joint, cellular life-Encompassing DIversity’ (*JEDI*) marker as a simple, effective and quantitative measure to assess and monitor biodiversity. The *JEDI* marker is a ribosomal RNA gene fragment that can be amplified from all domains of life using a single pair of PCR primers. We demonstrate the applicability and effectiveness of this approach for assessing biodiversity across ecological and biological scales, from holobionts to diverse ecosystems. In addition, we provide an automated bioinformatic workflow to support the standardised and reproducible analysis of the *JEDI* marker in future studies. While this approach is not free from trade-offs, we argue that its advantages outweigh its limitations by providing a unique, operational and scalable solution that builds on established infrastructure to integrate the microbial majority into biodiversity assessments and provide fundamental insights into organismal dynamics and associations across domains of life. Thus, the *JEDI* marker approach addresses the urgent need for a universal and standardised framework to effectively measure and monitor biodiversity at planetary scales in an era of profound global change.

## INTRODUCTION

Over the course of billions of years, evolution has driven the diversification and complexification of life, resulting in the rich diversity of organisms that inhabit today’s biosphere, which we collectively refer to as *biodiversity*. Beyond its intrinsic value, biodiversity forms the foundation of ecosystems’ function, health and habitability, and thus homeostasis of the Earth system^1–3^. Biodiversity also underpins ecosystem services that human societies depend on, from food production and water purification to the formation of soil and decomposition of waste, contributing an estimated $40 trillion per year—nearly half of the global Growth Domestic Product^4–6^. Yet, biodiversity is under severe threat from anthropogenic activities and climate change, with alarming rates of decline in plant and animal species and their associated genetic diversity^7–9^. Overall, the decline of biodiversity is recognised as one of the greatest existential threats facing humanity^10,11^. As such, the urgency to document, monitor, and understand changes in Earth’s biota has never been greater.

Despite its vital importance, biodiversity remains a highly contingent and inconsistently defined concept. Biodiversity can be perceived from different biological levels– genomic, metabolic, morphological, physiological or behavioural– with each providing distinct and often disconnected insights into biological complexity^12^. It can also be perceived across vastly different ecological scales, from the diversity of individual organisms, including their associated symbionts, to the collective diversity within an ecosystem or the entire biosphere. Biodiversity also encompasses the entire tree of life and organismal size spectrum, from single-celled prokaryotes and eukaryotes to colonial or multicellular organisms, such as metazoans and plants. Despite its breadth, biodiversity is typically perceived and measured through species. However, the difficulty of defining and delineating species consistently across the tree of life, introduces additional ambiguity that undermines the comparability and robustness of biodiversity assessments^13–15^. As a result of the multitude of perspectives and sheer scope of the concept, studying biodiversity has become inherently multidisciplinary, with disparate approaches across fields leading to a highly fragmented landscape of experts and knowledge. Over time, this fragmentation has fostered conceptual fluidity, resulting in inconsistencies in how biodiversity is defined, measured, and monitored across studies^15,16^. Therefore, while the biodiversity crisis is often considered as one of the greatest threats to humanity and the Earth’s biosphere, ironically, we still lack a cohesive and comprehensive understanding of biodiversity itself.

Current approaches used in biodiversity assessments to inform policy, society and guide conservation not only lack standardisation and suffer from scalability issues, but they also provide a severely biased and constrained perspective of life on Earth. Global biodiversity assessments thus far^17,18^ have primarily centred on enumerating species of macro-organisms, e.g. plants and animals, leading to estimates of up to eight million species on Earth^19^. However, this macro-organism-centric view overlooks the vast majority of life’s diversity, which is contained within microorganisms, such as archaea, bacteria, and protists^20^. This “microbiodiversity” has stood as the dominant form of life since its first emergence on Earth, diversifying over billions of years into a remarkable genomic, metabolic, and cellular diversity before giving rise to more complex multicellular organisms. In addition to representing the majority of Earth’s biodiversity, microorganisms have been instrumental in shaping the biosphere that we inhabit today and continue to form the foundation of food webs, drive biogeochemical cycles and serve essential roles as symbionts, without which macro-organisms could not exist^21–23^. The ecological significance of microorganisms has also been recently exemplified in the context of ecosystem recovery after perturbation or damage, with evidence that they can significantly improve the rate, resilience and overall success of ecosystem restoration efforts^24,25^. Integrating microorganisms into biodiversity assessments is thus of paramount importance, not only to gain a more complete and accurate perspective of biological diversity across taxonomic, ecological and environmental axes, but also to inform decisions on how to conserve and preserve natural ecosystems and effectively mitigate biodiversity loss. To address this need, new strategies must be developed that unify the assessment of biodiversity across the tree-of-life within a scalable and standardised framework.

In this study, we demonstrate that amplicon sequencing using the small subunit ribosomal RNA (SSU rRNA) gene primers 515F-Y/926R^26^ provides a robust and scalable approach for quantifying and analysing biodiversity across biological and ecological scales. Through *in-silico* analyses, we show that these primers provide an extensive coverage of organisms across the tree of life, comparable or superior to taxon-specific primer sets, while also effectively capturing the diversity of eukaryotic plastids. Based on these properties, we propose the SSU rRNA gene fragment amplified by these primers as the ‘Joint, cellular-life-Encompassing DIversity’ (*JEDI*) marker, and term the amplicon sequencing of these fragments as the *JEDI* marker approach. To support this, we demonstrate the applicability and effectiveness of the *JEDI* marker approach for quantifying and monitoring biodiversity across scales, from individual holobionts to whole ecosystems, including ocean water, sediments, and soils. We also develop a new bioinformatic function, the ‘*consensus merge pairs*’ approach, to ensure the standardised and reproducible processing of JEDI marker data in future studies. Together, our findings, conceptual considerations and practical implications provide strong support for adopting the *JEDI* marker approach as a unifying and practical framework to assess and monitor Earth’s biodiversity, addressing critical knowledge and methodological gaps and supporting a more holistic understanding of the patterns, processes, and drivers that shape life on our planet.

## RESULTS

### Universal primers capture a substantial fraction of diversity across the tree of life

Assessing biodiversity through DNA sequencing typically requires the use of primers that bind to conserved sites within genes, enabling amplification and sequencing of the variable regions between them. To be effective, such primers must capture a broad range of taxa, provide sufficient resolution to distinguish among them, and accurately reflect their relative abundances in the original environmental sample. While primers that target hypervariable regions of the SSU rRNA gene have been widely used to assess biodiversity, they typically target specific groups of organisms, such as taxa within Bacteria and Archaea or Eukaryota^27–29^. However, Parada et al, building on earlier efforts, developed a single primer set (515F-Y/926R) - hereon referred to as universal primers - capable of targeting the V4-V5 regions of the SSU rRNA gene across all domains of life. Thus, these primers present a unique opportunity to move beyond taxonomically and conceptually constrained and fragmented approaches toward a unified framework for global biodiversity assessments. While previous studies have demonstrated that the universal primers outperform taxon-specific primers in capturing diversity within ocean samples^29^, a comprehensive evaluation of the sequence diversity they capture across the tree of life and how this aligns with formally recognised taxonomic species, the foundational unit in biodiversity assessments, remains lacking.

To address this, we constructed curated reference datasets of prokaryotic and eukaryotic sequences to enable robust *in silico* comparisons of primers targeting different hypervariable regions of the SSU rRNA gene. Such curation is critical for minimising biases, as many of the sequences in reference databases do not span all hypervariable regions. These biases are particularly pronounced when comparing eukaryote-targeting V4 and V9 primers, where the V9 reverse primer target is frequently missing, leading to pronounced discrepancies in the diversity captured *in silico* compared to in environmental samples^29–31^. To minimise these biases, we aligned sequences from the PR2^32^ and SILVA^33^ databases to the backbone SSU rRNA gene alignment of SILVA and extracted those that spanned different primer binding positions (Fig. 1a). Through this, we devised a curated reference sequence (REF-SQ) dataset for the V3-to-V5 region, encompassing 17,872 archaeal and 364,970 bacterial sequences, and for the V4-to-V9 region, encompassing 66,885 eukaryotic sequences. Additionally, we applied a second curation step, retaining only those sequences representing validly named species (based on valid binomial nomenclature) to enable robust comparisons with taxonomic species diversity. The curated reference species (REF-SP) dataset of the V3-to-V5 region contains 535 archaeal and 14,303 bacterial species, while the V4-to-V9 REF-SP dataset contains 13,399 eukaryotic species.

**Figure 1.**
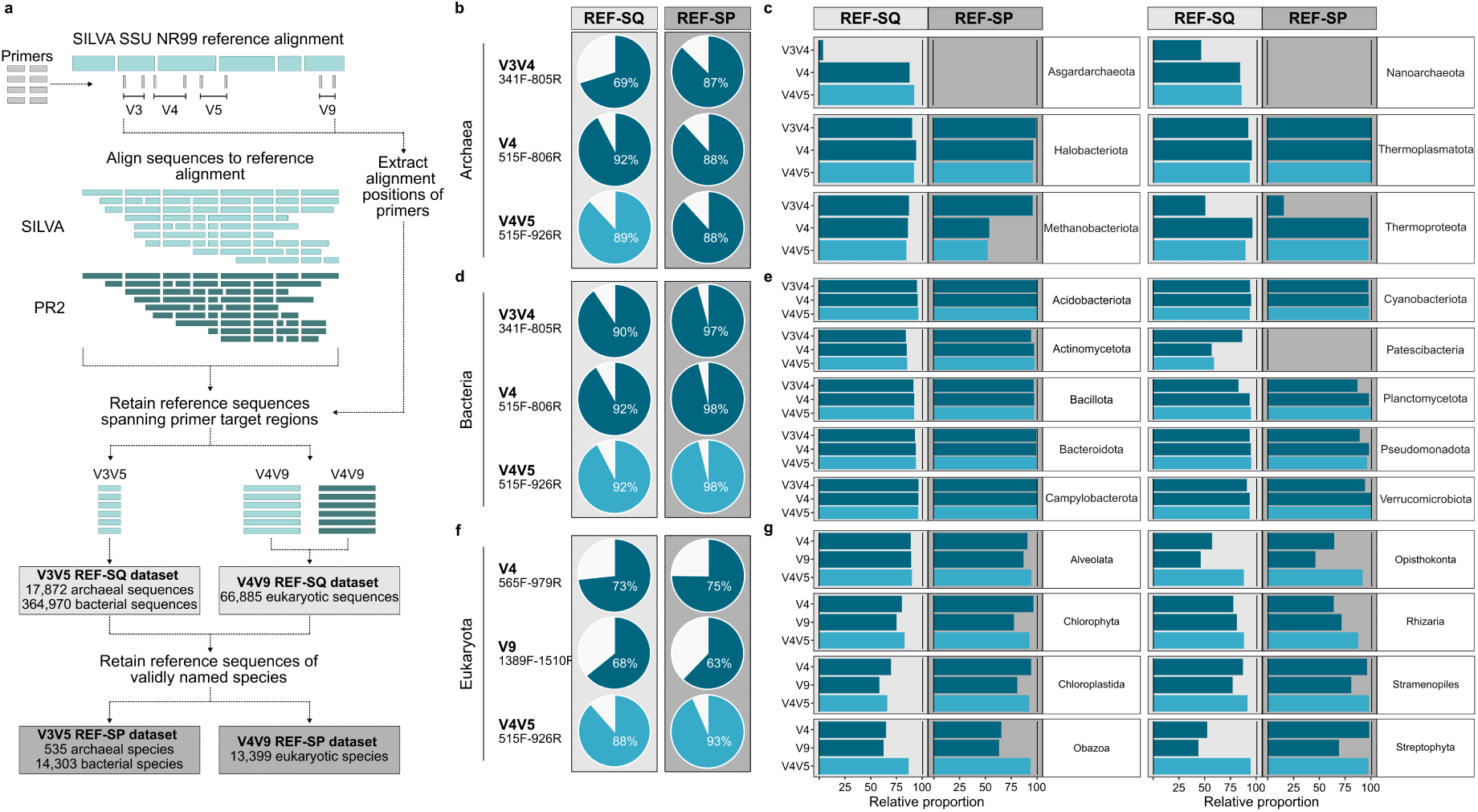
*In silico* assessment of universal and taxon-specific primers. **a)** Pipeline developed to generate curated datasets of SSU rRNA gene sequences spanning different variable regions of the SSU rRNA gene. The curated datasets were then used as a reference for a blastn search to assess the coverage of primers targeting differen t variable regions. **b)** Proportion of reference sequences (REF-SQ) and reference species (REF-SP) of Archaea captured by different primer pairs (allowing up to two mismatches). **c)** Primer coverage of major phyla of Archaea within the REF-SP and REF-SQ datasets, with empty panels representing taxa with no named species. **d)** Proportion of REF-SQ and REF-SP of Bacteria captured by different primer pairs (allowing up to two mismatches). **e)** Primer coverage of major phyla of Bacteria within the REF-SP and REF-SQ datasets, with empty panels representing taxa with no named species. **f)** Proportion of REF-SQ and REF-SP of Eukaryota captured by different primer pairs (allowing up to two mismatches). **g)** Primer coverage of major taxa of Eukaryota within the REF-SP and REF-SQ datasets.

Leveraging these curated reference datasets, we systematically assessed the extent of biodiversity captured *in silico* by the universal primers in comparison to those widely used for targeting prokaryotic and eukaryotic organisms separately (Table 1). The universal primers achieved an extensive coverage of prokaryotic diversity, capturing 92% of sequences and 96% of species within Bacteria and 89% of sequences and species within Archaea, comparable to the V4 and higher than the V3V4 prokaryotic-specific primers (Fig. 1b and 1d). While all primers demonstrated high coverage across major bacterial phyla, except for Patescibacteria and Spirochaetota (Fig. 1e and Supp. Fig. 1), clear differences were observed within Archaea. In particular, the universal and V4 primers exhibited a low coverage of species diversity within Methanobacteriota, while the V3V4 primers suffered from a low coverage of Asgardarchaeota, Nanoarchaeota and Thermoproteota (Fig. 1c). For eukaryotes, the universal primers exhibited the most extensive coverage of the primer sets tested, with 88% of sequences and 93% of species covered, followed by the V4 primers that captured 73% of sequences and 75% of species (Fig. 1f). The reduced coverage of eukaryote-targeting primers stems from taxon-specific biases (Fig. 1g and Supp. Fig. 1), including a low coverage of Obazoa, Opisthokonta and Streptophyta by V4 and V9 primers. We also observed that these differences in coverage were not influenced by the number of mismatches between the primers and reference sequences (Supp. Fig. 2). To provide an alternative perspective, we also compared the species-level coverage of the primers considering only forward primer matches, which revealed higher coverages for the V9 than the V4 primers (Supp. Fig. 3). These findings demonstrate that the universal primers provide an extensive coverage of organismal diversity across all three domains of life, which is greater than or comparable to the combined coverage of prokaryotic- and eukaryotic-specific primer sets.

**Table 1.**
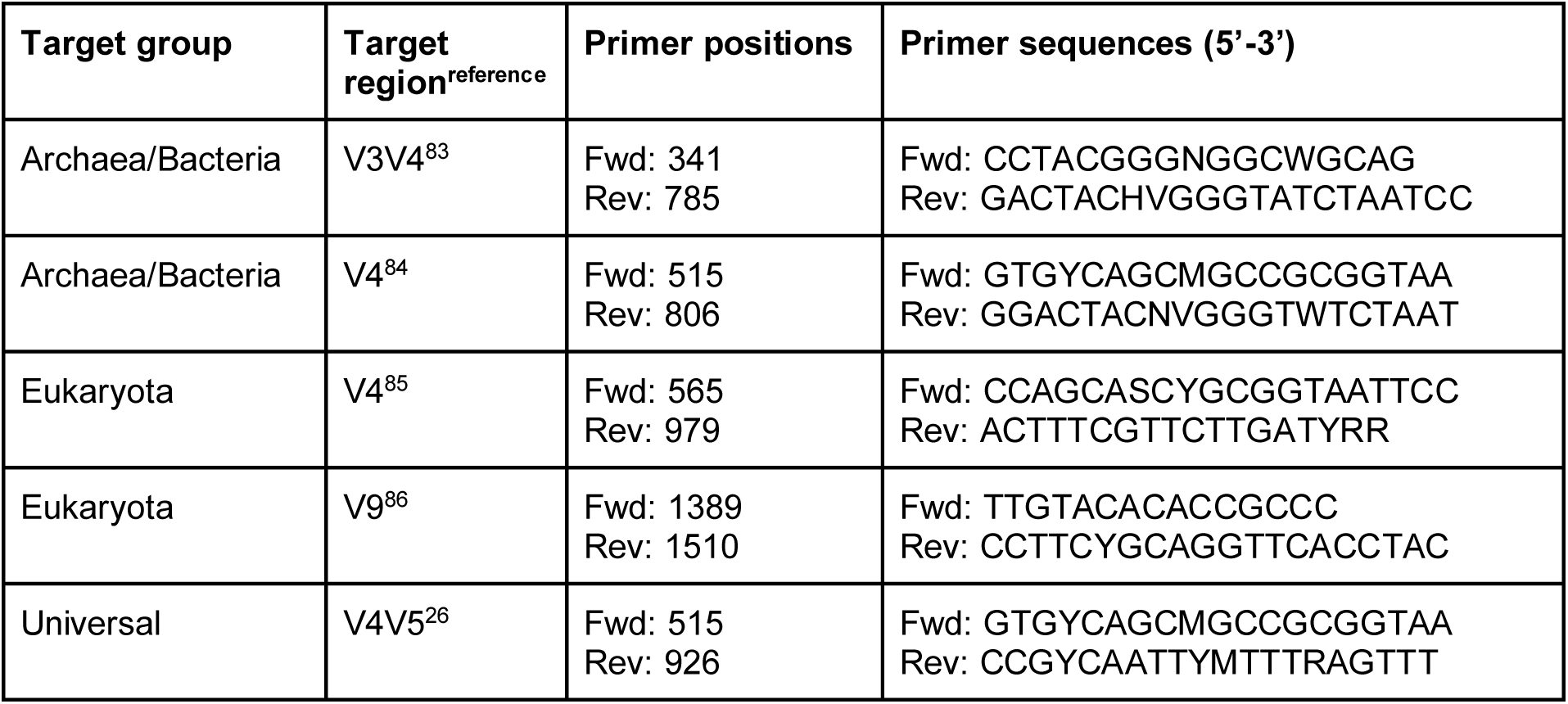
Primer pairs used for *in silico* comparisons. For prokaryotic-targeting primers, positions are based on *E. coli*, and for eukaryotic-targeting primers, positions are based on *S. cerevisiae*.

To further broaden and complement our assessment of the diversity captured by the universal primers, we explored their ability to target SSU rRNA genes encoded by eukaryotic organelles. To achieve this, we obtained complete mitochondrial (n = 18,206) and chloroplast (n = 13,342) genomes from the NCBI-organelle database and assessed the extent of diversity covered by the universal primers through a blast-based analysis (allowing up to two mismatches). We observed marked differences in the coverage of organelles, with only 4.8% of mitochondria but 99.6% of chloroplasts (sequences and species) captured by the universal primers. These findings suggest that the universal primers hold limited utility for profiling mitochondria but great potential for capturing chloroplast diversity and, by extension, the profiling of photosynthetic eukaryotes– further extending the breadth of biodiversity insights that can be achieved.

Given the universality and extensive coverage of diversity provided by the universal primers, we propose that the fragment of the rRNA gene they target can serve as a ‘Joint cellular-life-Encompassing DIversity’ (JEDI) marker, providing a unified and universal framework for assessing biodiversity.

### Employing the JEDI marker for quantifying biodiversity

Having demonstrated the universality of the JEDI marker, we next assessed the resolution it provides for distinguishing between species and thus aligning with current biodiversity assessments. In DNA-based surveys of biodiversity using the SSU rRNA gene, the amplified and sequenced regions - known as amplicons - are typically processed into Amplicon Sequence Variants (ASVs)^34^. Defined as representing unique sequences, ASVs have become the standard unit of diversity in amplicon-based studies because they are simple, stable, reproducible, and comparable across samples if processed through a standardised workflow^34^. However, ASVs do not always align with taxonomic species. In some cases, they provide a finer-grained resolution^35^, with multiple ASVs for a single species, while, in other cases, multiple species are encapsulated under a single ASV^36^. This variability poses challenges when aligning molecular- and species-based biodiversity data. While species units may arguably not be an accurate representation of ecological diversity and cannot be defined on common criteria across the tree of life, they remain foundational in classification schemes and still serve as the primary currency of conservation efforts and policy frameworks. Therefore, understanding how *JEDI* marker-based ASVs relate to species units is essential for integrating DNA-based biodiversity surveys with species-based frameworks and preserving continuity with historical records.

Prior to assessing the resolution of *JEDI* marker ASVs, we first addressed a technical challenge: classical pipelines for inferring ASVs are ill-suited for the *JEDI* marker due to variability in its length across taxonomic groups. In particular, the elongation of the V4 region in eukaryotes results in amplicons that exceed the length coverable by current paired-end sequencing technologies (Fig. 2a), causing the loss of eukaryotic reads during the overlap-based merging step. Existing solutions involve either retaining only one read or using a taxonomic-informed approach to identify and merge reads of 16S rRNA genes and concatenate those of 18S rRNA genes^37,38^. However, taxonomic pre-sorting can introduce classification bias and may enforce concatenation or merging unnecessarily - for example, some eukaryotic sequences (e.g. within Microsporidia) are short enough to be fully covered by paired-end reads, while some bacterial JEDI markers exceed the maximum length for merging (Fig 2a). To overcome these limitations, we developed a new strategy—termed the *consensus merge pairs* approach—that has been integrated into the nf-core/ampliseq^39^ workflow since version 2.13^40^. This method merges overlapping paired-end reads and concatenates non-overlapping ones in a single step, without pre-partitioning based on taxonomy (see Methods) (Fig. 2b). While inspired by the work of Needham and Fuhrman^41^, our approach incorporates optimised alignment scores and adaptive overlap thresholds to ensure that concatenation only occurs for truly non-overlapping reads, avoiding artefacts from mismatches or sequencing errors in overlapping regions (see Supplementary Information). The *consensus merge pairs* approach thus ensures the accurate and scalable inference of ASVs from *JEDI* marker amplicons across all domains of life.

**Figure 2.**
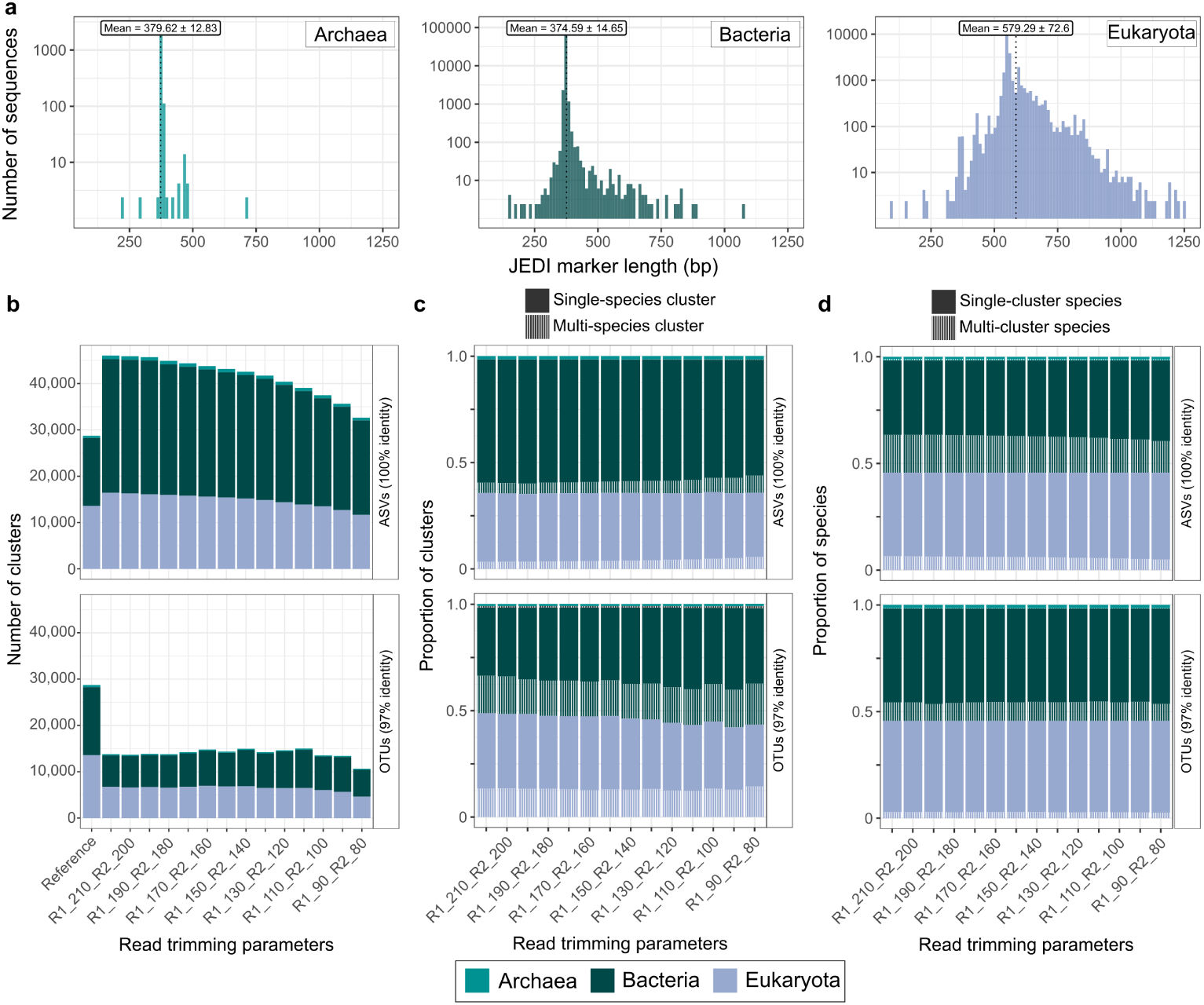
Evaluation of *JEDI* ASVs as a proxy for taxonomic species diversity. **a)** Distribution of *JEDI* marker lengths across Archaea, Bacteria and Eukaryota reference species. **b)** Comparison of the number of reference species and the number of clusters obtained through the *consensus merge pairs* approach using both a 100% and 97% global-alignment identity threshold with cd-hit and different read length trimming parameters. **c)** Proportion of clusters formed at 100% sequence identity, using different read length trimming parameters, which contained a single or multiple species. **d)** Proportion of taxonomic species contained within a single cluster or split across multiple clusters with 100% sequence identity across different read length trimming parameters.

To explore how *JEDI* marker ASVs relate to taxonomic species units, we extracted and analysed the respective regions from the REF-SP dataset. For this, we simulated paired-end amplicon sequencing by extracting the *JEDI* marker from 27,476 reference species matched by the universal primers (allowing two mismatches) and splitting these regions at varying lengths to reflect reads post quality trimming. By applying a merging and concatenation approach analogous to *consensus merge pairs*, we recovered 32,010-46,037 ASVs depending on the paired-end read lengths used (Fig. 2b). These numbers suggest that *JEDI* marker ASVs may overestimate species-level diversity by 11–58%. However, these inflated values are largely attributed to Bacteria, with up to 96% more ASVs than reference species compared to only 29% more ASVs for Eukaryota (Fig. 2c) when using the longest read lengths. To understand how this overestimation arises, we explored the distribution of species within and across ASVs. From the perspective of species information loss, we found that only 6% of archaeal, 8% of bacterial, and 10% of eukaryotic taxonomic species were grouped into the same ASV, indicating low rates of under-splitting (Fig. 2c). From the perspective of inflating diversity estimates, we observed that most species were represented by a single ASV, encompassing 56% of species for Archaea, 66% for Bacteria, and 86% for Eukaryota, while the remainder were split across multiple ASVs (Fig. 2d). For comparison, clustering the *JEDI* marker into Operational Taxonomic Units (OTUs) at 97% identity– a threshold proposed to better align with species boundaries–resulted in a 55-58% underestimation of diversity due to the merging of multiple species into single clusters, affecting 44% of archaeal, 36% of bacterial and 28% of eukaryotic species (Fig. 2c). Together, these findings demonstrate that, while *JEDI* marker ASVs typically capture a finer-grained resolution than species units, they rarely lose species-level information.

Based on these observations, we propose that amplicon sequencing of the *JEDI* marker– which we refer to as the *JEDI* marker approach –can enable a unique, objective and quantifiable, tree-of-life-scale measure of biodiversity. This measure is underpinned by unique variants (i.e. ASVs) of the *JEDI* marker as the common denominator. To ensure robust and stable inference of *JEDI* marker ASVs, and thus of biodiversity estimates, we provide the automated ‘*consensus merge pairs*’ approach. While these estimates will provide a finer-grained perspective than those based on taxonomic species, the comprehensive *in silico* comparison performed here provides quantitative insights into the discrepancy between the two, which can serve to support the future integration of biodiversity measurements from existing species-based approaches and those of the *JEDI* marker.

To demonstrate the effectiveness and applicability of the *JEDI* marker approach and support our proposal, we applied the universal primers in four different projects to assess biodiversity across different biological and ecological scales, from holobionts to across different ecosystems.

### Application example 1: Tracking cross-domain diversity and dynamics within single-cell and multi-cellular holobionts

It has become increasingly evident that the ecology and evolution of all organisms is highly influenced by their associated microbiomes. The realisation of this interdependence has led to the *holobiont* concept^42^, which considers an organism and its associated microbiome as a single biological unit. Host-associated microbiomes play essential roles in the growth, development, physiology, ecological fitness, adaptation and evolution of their hosts, influencing processes such as nutrient acquisition, immune modulation, stress tolerance, and disease resistance^43–45^. While amplicon sequencing is a powerful technique for gaining insights into the diversity and dynamics of host-associated microbiomes, the use of taxon-specific primers has hampered our understanding of interactions across domains and their dynamics in relation to the host. Here, we demonstrate how the *JEDI* marker can help address these knowledge gaps by applying it to a unicellular and multicellular holobiont system.

To investigate the microbiome associated with single cells of diatoms in coastal waters, we isolated 94 *Rhizosolenia* sp. cells through image-based cell sorting into a 96-well plate (Fig. 3b). Application of the *JEDI* marker approach to wells containing single-cells resulted in the recovery of 162 ASVs spanning Bacteria (n=148), Eukaryota (n=6), eukaryotic plastids (n=5), and Archaea (n=3). The recovery of nuclear, plastidial and mitochondrial ASVs classified as *Rhizosolenia* from 99%, 100% and 98% of sorted cells, respectively, confirmed sorting accuracy (Fig. 3a). Alongside these host-derived signals, we also detected co-occurring ASVs in 98% of cases, with a mean of eight ASVs associated with each host cell (Fig. 3a).

**Figure 3.**
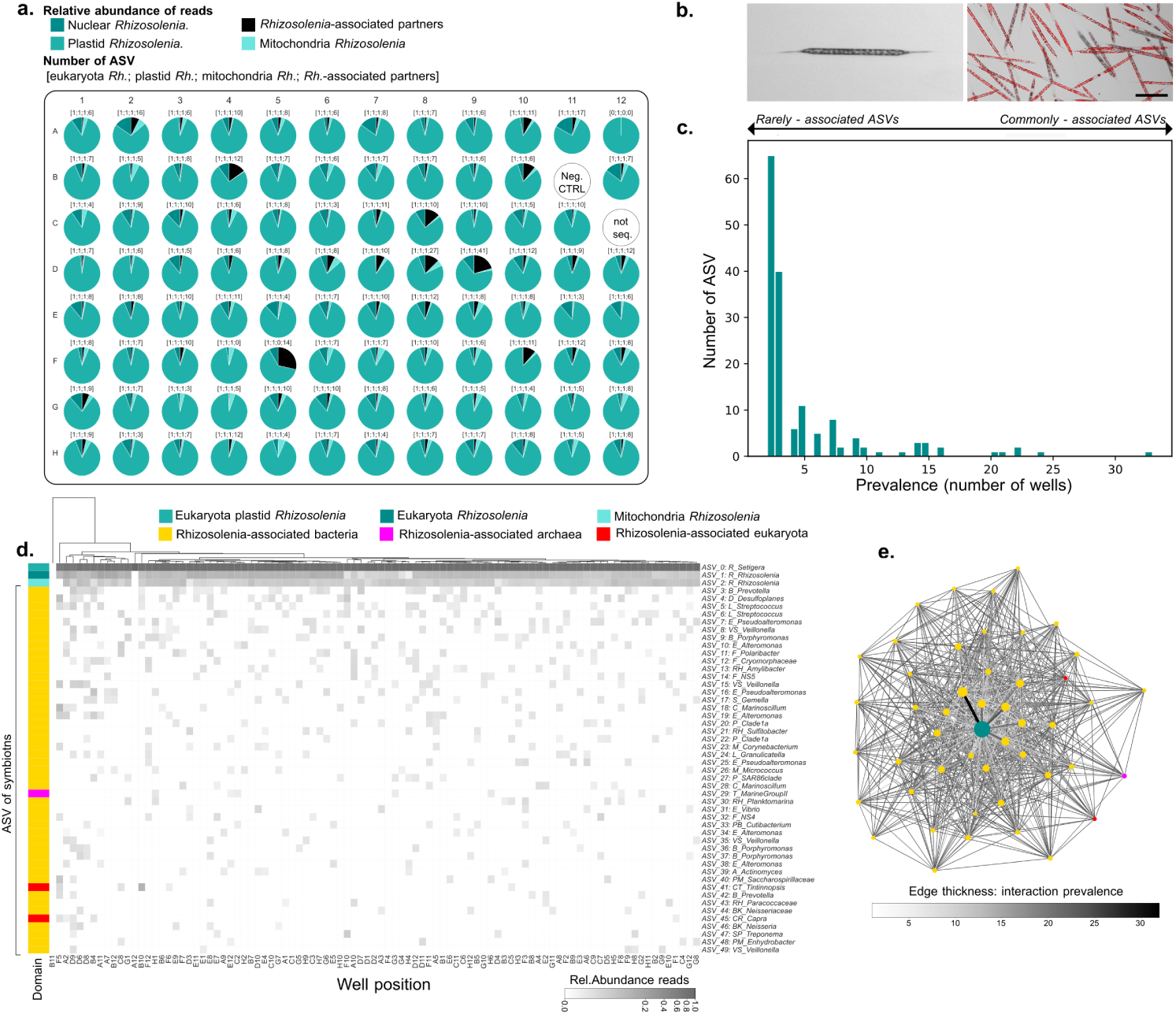
Profiling unicellular holobionts with the JEDI marker. **a)** Pie chart of the relative abundance of JEDI marker Amplicon Sequence Variants in each individual well where a single cell of *Rhizosolenia* was automatically sorted; *Rhizosolenia* nuclear 18S (dark green), *Rhizosolenia* plastidial 16S (green), *Rhizosolenia* mitochondria (light green), other associated microorganisms (black, including Bacteria, Archaea, Eukaryota and Eukaryota plastid). C12 was not sequenced, B11 is the negative control with no *Rhizosolenia* . **b)** Left: image of *Rhizosolenia* from the image-enabled cell sorter used to produce the single-cell plate; right: confocal microscopy image of *Rhizosolenia* cultured after isolation (red: chlorophyll autofluorescence), scale 100µm**. c)** Frequency distribution of the symbiotic ASVs prevalence across wells. **d)** Heat map of the relative abundance of the 50 most prevalent ASVs. The wells (holobionts) are clustered by similarity, the ASV are ordered from the most to the least prevalent. Order abbreviation: R : *Rhizosolenia*, RK: Rickettsiales, B: Bacteroidales, D: Desulfovibrionales, L: Lactobacillales, E: Enterobacterales, VS: Veillonellales-Selenomonadales, F: Flavobacteriales, RH: Rhodobacterales, S: Staphylococcales, C: Cytophagales, P: Pelagibacterales, M: Mycobacteriaceae, MC: Micrococcales, PM: Pseudomonadales, T: (family) Thermoplasmata, PB: Propionibacteriaceae, A: Actinomycetales, CT: Choreotrichida-Tintinnida, BK: Burkholderiales, CR: Craniata, SP: Spirochaetales. **e)** Realised co-occurrence network. Each node represents an ASV and each edge corresponds to a physical interaction. Node color indicates taxonomy as in **d)**, node size indicates prevalence of the ASV across 95 wells and edge thickness indicates prevalence of the interaction across 95 wells.

To explore the structure and composition of diversity associated with cells of *Rhizosolenia* sp., we assessed the prevalence and co-occurence of ASVs across wells (Fig. 3c). We identified both commonly detected ASVs, comprising members of *Pseudoalteromonas*, *Alteromonas*, *Porphyomonas,* and *Polaribacter,* and rarely detected ASVs, including *Tintinnopsis* ciliates (n=5), *Thraustochytriaceae* protists (n=3), photosynthetic eukaryotes such as *Bathycoccus* (n=2), and microbial taxa like *Vibrio splendidus* and *Thermoplasmatota* (Fig. 3d). While the composition identified with each cell was dominated by the host signal (96.6±4.3%reads), the rarely detected ASVs typically represented a larger fraction of the community than the commonly detected ASVs, particularly *Tintinnopsis* (2.29±4.25% reads), Saccharospirillaceae (0.9±1.45% reads) and *Marinoscillum* (0.87±2.03% reads). Co-occurrence network analysis (Fig. 3e) further revealed structured associations within the microbiome, including a potential central role for *Cryomorphaceae* and *Porphyromonas* (each with 43 co-occurring partners), as well as consistent interactions among marine bacteria like *Pseudoalteromonas*, *Porphyromonas*, and *Polaribacter*, suggesting potential interactions within the holobiont. Together, these observations provide insights into the extent and composition of diversity associated with single diatom cells in coastal ocean waters, while demonstrating that the *JEDI* marker approach, when combined with cell-sorting techniques, can facilitate the simultaneous analysis of both the host and its associated microbiome on a single-cell level.

Building on its utility for profiling unicellular holobionts, we next applied the *JEDI* marker approach in a year-long survey to investigate the dynamics of a multicellular holobiont, the brown macroalga *Ascophyllum nodosum*, a key habitat-forming species of intertidal zones of the North Atlantic^46^. Brown macroalgae are known to harbour rich microbiomes including epi- and endosymbiotic prokaryotes and eukaryotes^47–49^, with the *A. nodosum* holobiont also containing two fungal members, *Mycophycias ascophylli* and *Moheitospora* sp.^50^.

Through the application of the *JEDI* marker approach to algal tissue samples collected monthly between November 2021 to November 2022, we detected a rich diversity of microorganisms, comprising 1430 - 3197 ASVs, predominantly attributed to bacteria (Fig. 4a). The dynamics of the microbiome in relation to the host– a unique aspect enabled by universal primers– varied over the annual cycle, with a reduction in the microbiome signal in early spring (Fig. 4b), which coincides with the known period of annual epidermal shedding of the algae^51^. We further disentangled individual temporal dynamics between the bacterial, eukaryotic and fungal fractions of the microbiome, suggesting different drivers or controlling forces (Fig. 4c). To assess potential interactions or co-regulated members of the microbiome, we leveraged the host signal as a reference for normalisation and compared the dynamics between populations (ASVs). Through this, we identified strong positive correlations (correlation coefficient >0.5, adjusted p-value <0.05) between the dynamics of 41 bacterial populations and the two fungal partners, including 9 bacterial populations with *Moheitospora* sp. and 37 bacterial populations with *Mycophycias* sp. (Fig. 4d). Among those bacterial populations with the highest correlation coefficients were members of *Pirellulaceae* (Planctomycetes), *Rhodobacteraceae* (Alphaproteobacteria), *Saprospiraceae* (Bacteroidia) and *Sphingomonadaceae* (Alphaproteobacteria). Further supporting the unique insights gained by the *JEDI* marker approach, these correlated changes in relative abundance were unobservable when comparing the signal of the bacterial populations to that of the fungi obtained using an independent ITS marker, i.e. without normalisation by the host signal. These findings highlight the dynamic nature of macroalgal-associated microbiomes and the unique insights on cross-domain dynamics within holobionts that can be facilitated by the *JEDI* marker approach.

**Figure 4.**
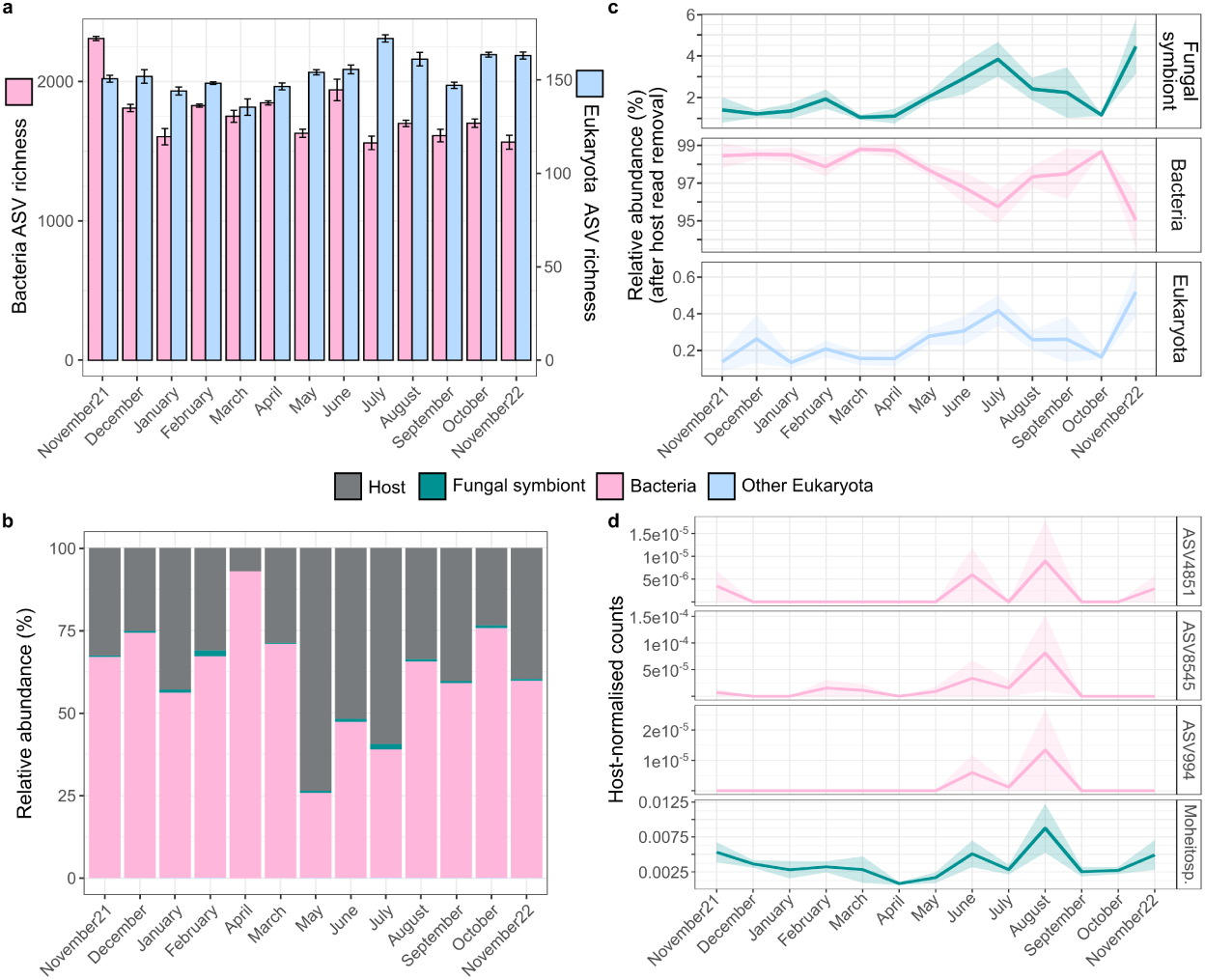
Unveiling of macroalgal host-associated microbiome diversity and dynamics through the JEDI marker approach. Tissue samples from the brown seaweed *Ascophyllum nodosum* were collected from six individuals within the same location, once per month over the course of an annual cycle. **a)** Mean and standard error of bacterial and eukaryotic ASV richness across the macroalgal samples determined through 100 iterations of subsampling and estimating richness. **b)** Relative composition of the holobiont. **c)** Relative abundance dynamics of Bacteria, Fungi and other eukaryotes after removal of host reads. Based on the taxonomic classifications, ASVs were grouped into three categories, representing the known fungal symbionts and the remaining bacterial and eukaryotic ASVs. Lines represent the mean and clouds the standard error of the mean relative abundance observed across the six algal samples collected during each month. **d)** Pairwise Pearson’s correlation analysis was performed between each ASV and those of the two fungal symbionts (*Moheitospora sp.* and *Mycophycias ascophylli)*. Strong correlation coefficients with a value of >0.7 and an adjusted p-value <0.05 were retained. The most abundant ASVs exhibiting a correlation with Moheitospora were used for visualisation.

### Application example 2: Tracking multi-year, cross-domain diversity dynamics in an ocean ecosystem

The application of amplicon sequencing in marine time-series studies– which involve frequent and periodic sampling at a fixed location –has been fundamental for uncovering the composition and dynamics of prokaryotic communities over daily, seasonal and annual timescales^52–54^. More recently, it has also been extended to track eukaryotic communities over time within ecosystems, revealing distinct temporal fluctuations shaped by different environmental drivers along with biotic interactions^38,55,56^. However, in most studies, prokaryotic and eukaryotic communities have been studied using independent primers. As a result, we have a limited understanding of how diversity is structured at the tree-of-life scale and how ecosystems are ultimately shaped by dynamics and interactions that take place within and across domains of life. The *JEDI* marker approach presents a unique opportunity to address this and enable a more comprehensive and holistic perspective on diversity and community dynamics within ecosystems.

To explore the structuring of diversity and cross-domain dynamics over intra- and inter-annual scales in a marine ecosystem, we applied the *JEDI* marker approach to surface water samples collected over four years at the Roscoff time-series station in the Western English Channel^55^. Bi-weekly collection of plankton samples from two organismal size fractions (0.2 -3 µm and >3 µm) yielded a rich diversity of organisms, comprising 2,802 bacterial, 116 archaeal and 1,869 eukaryotic ASVs, despite relatively shallow sequencing depth (17,449±277 paired-end reads/sample). The sampled communities were consistently dominated by bacteria across all time points, accounting for 79.9 ± 8.6% of ASVs and 82.8 ± 7.6% of reads (Fig. 5b and 5c). However, we observed clear seasonal shifts in the structuring of diversity, with contrasting patterns between prokaryotes and eukaryotes. Notably, eukaryotic representation increased markedly during spring and summer in terms of both diversity and composition, comprising 20.6-23% of ASVs and 9.6-11.9% of reads, compared to 5.9-8.5% of ASVs and 0.7-1.6% of reads in winter– a three- to tenfold seasonal fluctuation. While similar seasonal trends have been reported from ocean time-series using taxon-specific primers^56^, the *JEDI* marker approach uniquely enables the simultaneous capture and comparison of cross -domain dynamics, revealing proportional shifts and structuring of community diversity and composition^38,41^.

**Figure 5.**
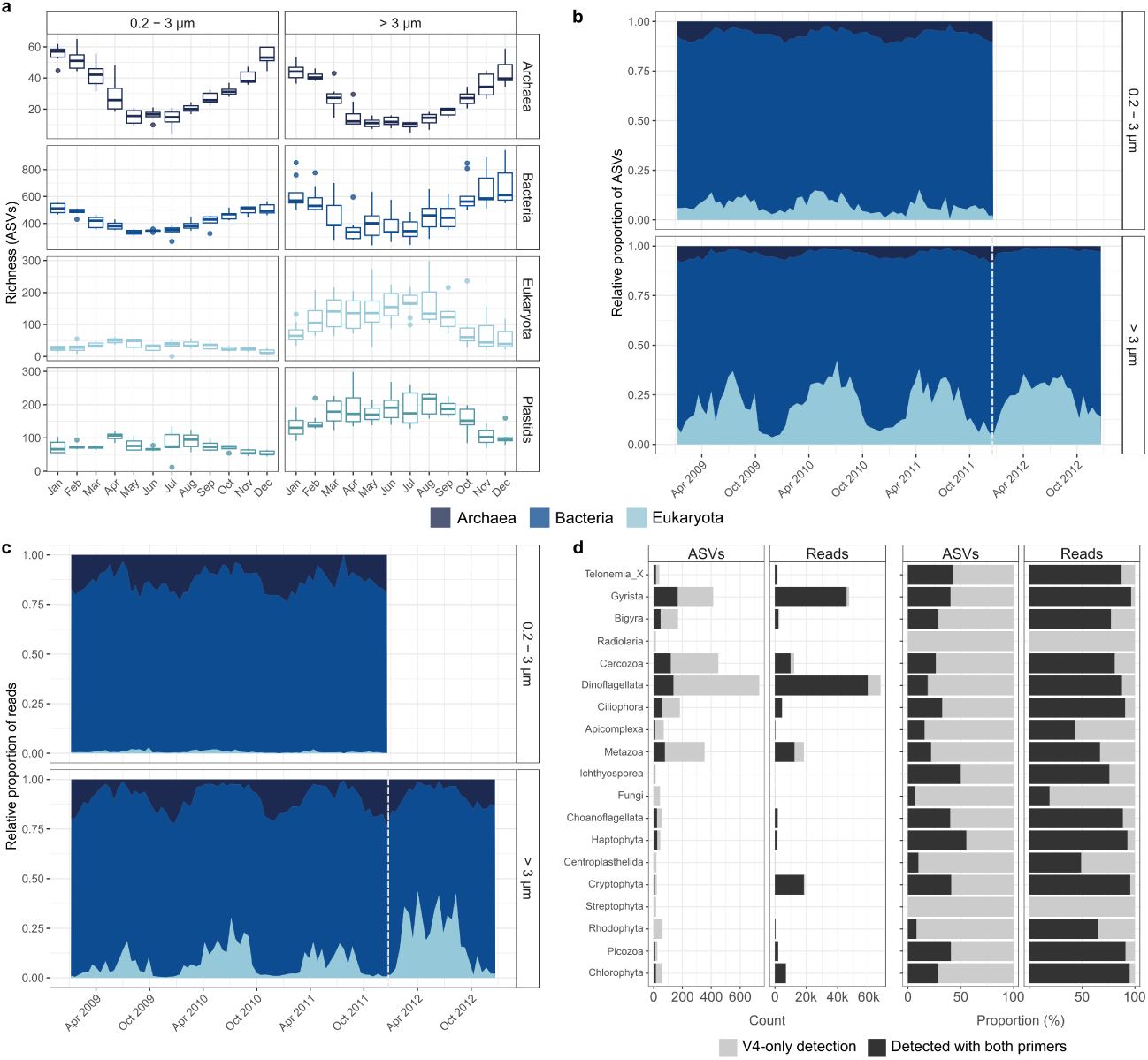
Cross-domain diversity and dynamics over four annual cycles at the Roscoff plankton time-series. Surface ocean samples were collected at biweekly intervals between 2009-2013 (2009-2011 for 0.2-3 µm fraction and 2009-2012 for >3 µm fraction) off the coast of Roscoff, as part of the SOMLIT-Astan time-series, and sequentially filtered through a 3 and 0.2 µm pore-size filter. The *JEDI* marker was amplified from both size fractions using the universal primers. **a)** Per-sample mean Richness (number of unique ASVs) of Archaea, Bacteria, Eukaryota and eukaryotic plastids recovered using the universal primers, after subsampling to the lowest count observed across the time-series. Samples are grouped by months. Temporal dynamics in the **b)** proportion of *JEDI* marker ASVs and **c)** proportion of reads attributed to Archaea, Bacteria and Eukaryota. Samples to the right of the dashed vertical line were those sequenced to a fourfold higher depth. **d)** Comparison of the number and proportion of ASVs and reads recovered for the major eukaryotic taxa by the V4 and universal primers from the >3 µm fraction samples across the entire time-series based on alignments of the shared regions (*i.e.* position 5-230 for the SSU V4V5 and position 1-226 for the 18S V4).

We next sought to evaluate the extent and dynamics of eukaryotic diversity observed with the universal primers in comparison to a taxon-specific primer set. To achieve this, we analysed an amplicon dataset generated from the eukaryote-targeting V4 primers applied to the >3 µm fraction samples of the same time-series. The V4 primers recovered a greater richness of eukaryotic ASVs (n=4,858) than the universal primers (n=1,689) (after subsampling to 9,944 reads per sample). Comparing the overlapping regions of the ASVs, we observed that JEDI marker ASVs were recovered by the V4 primers, representing only 20% of the total V4 ASVs. However, these shared ASVs accounted for 85% of sequenced reads (Fig. 5d), indicating that the *JEDI* marker approach effectively captured the most abundant eukaryotes across the time-series. To determine whether the lower richness observed with the *JEDI* marker approach was a consequence of the shallow sequencing depth, we compared samples from the first three years to those from the final year, during which sequencing depth in the >3 µm fraction was increased more than fourfold (from 17,449 ±277 to 63,855 ±1,491 paired-end reads). The greater sequencing depth resulted in a marked increase in the relative abundance of eukaryotes (from 3.2 ± 0.3% to 10 ± 1.3%) and yielded a diversity more comparable to that recovered with the V4 primers, evidenced through a reduced Jaccard distance (Supp. Fig. 4). These results demonstrate that the *JEDI* marker approach enables the detection and tracking of major eukaryotic populations over time in an ocean ecosystem, comparable to taxon-specific primers if amplicons are sequenced with sufficient depth.

### Application example 3: Measuring biodiversity across the land-sea interface

Studies employing amplicon sequencing to measure and track biodiversity have primarily been centered on single ecosystems, thus we have a limited understanding of how the composition, extent, and structure of biodiversity varies across ecosystems and at a global scale^57^. While feasibility was once a major obstacle to performing such assessments, this has been overcome through the reduction in cost and increase in throughput of DNA sequencing over the past decades. Therefore, we sought to investigate the suitability of the *JEDI* marker approach for facilitating integrated assessments of biodiversity across different ecosystems. To achieve this, we applied the *JEDI* marker approach with a standardised PCR and sequencing protocol to 23 samples collected across the land-sea interface, from soils, sediments and shoreline and coastal ocean waters.

We recovered a total of 94,115 ASVs, of which 92% represent Bacteria, 2.3% Archaea and 5.7% Eukaryota (of which 0.8% are plastidial). Rarefaction analysis revealed that the extensive sequencing depth employed, with an average of 1,770,262 ± 23,233 reads per sample, provided a substantial coverage of the biodiversity within the sampled communities (Fig. 6a). Given that such sequencing depths are often unachievable in large projects, we leveraged these rarefaction curves to estimate the extent of ASV diversity that could be captured across a range of shallower sequencing depths; 100,000 reads per sample would have captured 57-71%, while 250,000 reads per sample would have captured 74-87% of ASV diversity.

**Figure 6.**
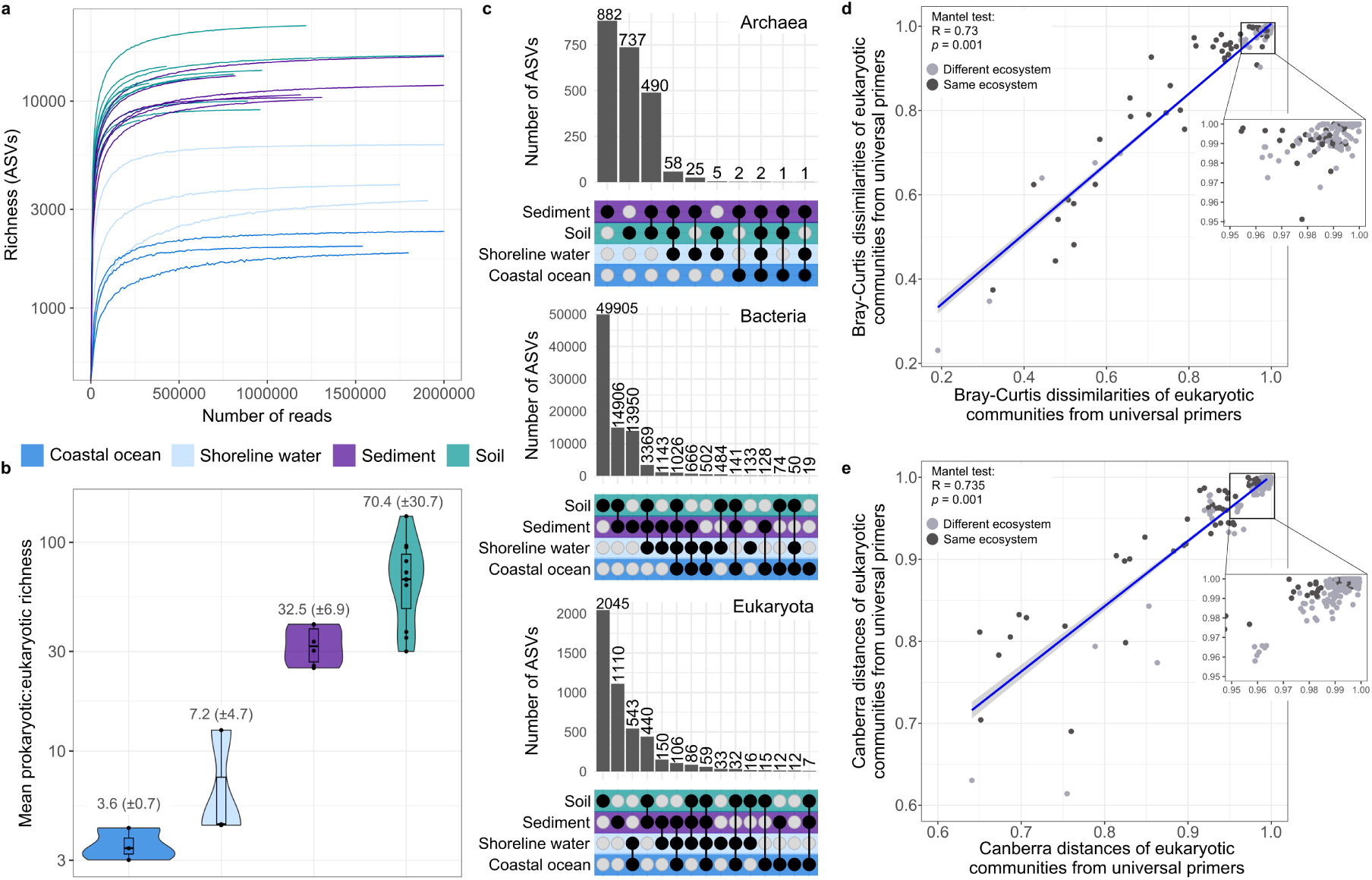
Alpha and beta diversity of soil, sediment, beach water and coastal ocean samples recovered through amplicon sequencing of the *JEDI* marker. a) Rarefaction curves of per-sample ASV richness determined through iterative subsampling of reads at 250 count intervals from 0 to 2,000,000. **b)** Mean per-sample prokaryotic:eukaryotic ASV richness determined through 100 iterations of subsampling and calculating richness. **c)** Upset plots illustrating the number of unique and shared ASVs across ecosystems. Comparison of mean between-sample Bray-Curtis dissimilarities **(d)** and Canberra distances **(e)** of eukaryotic communities recovered using the universal and eukaryote-targeting V9 primers. Dissimilarity/distance values were determined after subsampling to the lowest count observed in the universal primer dataset followed by exclusion of ASVs not assigned to Eukaryota.

We next compared the diversity and composition of ASVs detected within and across ecosystems. The richness of ASVs was highest in soils (n=11,420 ± 2835) and sediments (n=9268 ± 1630), which harboured threefold higher diversity than shoreline waters (n=3292 ± 1669) and sixfold higher diversity than coastal ocean samples (n=1512 ± 320) (Kruskal-Wallis; *H*=14.662, *p*=0.0021; Supp. Fig. 5). However, these community-level distinctions were primarily underpinned by Bacteria. In contrast, archaeal richness peaked in sediments (n = 231 ± 64) while eukaryotic richness was highest in shoreline waters (n=389 ± 91) and lowest in soils (n=185 ± 80) (Supp. Fig. 5). Consequently, the ratio of prokaryotic to eukaryotic ASV richness varied substantially across ecosystems, ranging from 3.5:1 in coastal ocean waters to 70:1 in soils (Fig. 6b). In addition to differences in the structuring of diversity at the domain level, we also observed distinct community compositions across ecosystems (Supp. Fig. 5), which were largely driven by ecosystem-specific ASVs. Ecosystem-specific ASVs were particularly rich in soils and sediments, with 52,799 and 16,076 unique ASVs, respectively (Fig. 6c). However, we also detected varying degrees of overlaps in diversity between ecosystems. For Bacteria and Archaea, the largest overlap in diversity was observed between sediments and soils followed by sediments and shoreline water, while for Eukaryota, the largest overlap was observed between beach water and coastal ocean (Fig. 6c). These findings highlight the distinct structuring of biodiversity across the land-sea interface, with unique insights into cross-domain patterns that are unattainable through taxon-specific primer sets.

We finally assessed whether the composition of diversity recovered with the *JEDI* marker approach is comparable to that detectable with a taxon-specific primer set. For this, we sequenced the same samples using a eukaryote-targeting V9 primer pair and compared the compositional differences detectable at equal sequencing depths (subsampling to the lowest read count across both datasets). Focusing only on the eukaryotic fraction of communities, the compositional differences observed with the universal and the V9 primers were largely concordant (Mantel test; *R*=0.73, *p*=0.001) (Figure 6d). However, given that Bray-Curtis dissimilarities are biased towards abundant ASVs, we also compared compositional differences through Canberra distances, which revealed a comparable degree of concordance between the two primer sets (Mantel test; *R*=0.75, *p*=0.001) (Fig. 6e). Therefore, although taxon-specific primers may result in the recovery of a greater diversity of ASVs than the *JEDI* marker approach at equal sequencing depths, they provide a comparable resolution in the detection of compositional shifts in biodiversity across ecosystems.

## DISCUSSION

We are living in an era of profound global change that is placing immense pressure on Earth’s biota, with severe consequences for biodiversity and major ramifications for the functioning and homeostasis of ecosystems and human society^9,25^. However, our ability to assess, monitor, and mitigate these changes remains constrained by fragmented, species-centric, and macroorganism-biased approaches that overlook much of life’s diversity. In this study, we set out to evaluate a standardised and scalable approach that enables comprehensive assessment and quantification of biodiversity across the full spectrum of life, integrating both micro- and macro-biota, under a unified metric.

Amplicon sequencing of the JEDI marker offers a powerful and pragmatic solution to overcome these limitations while capitalising on existing data and established infrastructure. Amplicon sequencing is already the most widely employed technique for studying the diversity of microorganisms in the environment, with over 1.6 million samples analysed to date (https://microbeatlas.org/). Its widespread adoption in eDNA surveys has also demonstrated its utility and effectiveness for assessing the diversity of larger, multicellular organisms, from corals^58^ to fish^59^ and cetaceans^60^. Supporting this has been technological advancements and cost reductions in DNA sequencing, which allow for the parallel processing of hundreds of samples at a cost of tens of US dollars each, making large-scale monitoring feasible. In addition, amplicon sequencing is supported by mature and robust computational tools and standardised workflows that enable the reproducible and comparable analysis of sequence data. The *JEDI* marker approach thus builds on this mature foundation and is further supported by our development of the *consensus merge pairs* approach (integrated into nf-core/ampliseq), which enables automated, standardised processing of *JEDI* marker data into taxonomically annotated ASVs, thereby ensuring reproducibility and comparability of biodiversity assessments moving forward.

Using *JEDI* marker ASVs as the unit of measure for biodiversity is operationally practical and objective, and circumvents dependence on classification schemes and ambiguous species units^13^. Traditional classification schemes have been instrumental for organising the relationships between the observed life forms on Earth, but their reliance on species as the fundamental unit presents significant challenges. The concept of species is inherently ambiguous, with no universally consistent framework for delineation. While multiple species concepts have been proposed—such as the Biological Species Concept, which defines species based on reproductive isolation—they often lack applicability for asexual organisms and are ill-suited to unicellular eukaryotes^61^ and prokaryotes^62^. More recently, genomic and genetic-based approaches have emerged as powerful alternatives for discriminating species, particularly for microorganisms^63–65^. However, while these methods have advanced our understanding of diversification and evolutionary processes, they remain cost- and resource-intensive, and are constrained by our ability to reconstruct genomes of environmental populations–which is particularly challenging for eukaryotes with large and complex genomes^66^. Moreover, genomic insights have further revealed the limitations of species-centric frameworks in capturing ecologically meaningful diversity in natural communities, with individual species often encompassing a wealth of microdiversity structured into ecologically discrete and evolutionarily cohesive populations^67–69^. For these reasons, we advocate moving beyond species-based perspectives and adopting ASVs as an operational unit of diversity. While we recognise that ASVs vary in resolution across taxonomic groups, they nonetheless provide a consistent and reproducible unit derived from exact sequence information. Therefore, they offer a unified and universal metric that is not contingent on ill-defined units or unavailable taxonomic classifications, and, as demonstrated in this study, enable stable and reproducible estimates of biodiversity that can be compared across samples and studies.

Studies investigating the organismal make-up of ecosystems have, so far, primarily focused on specific taxonomic domains or trophic levels, providing valuable but highly fragmented perspectives. Such targeted approaches constrain our ability to understand how biodiversity is structured across the entire tree of life and to uncover associations that span domains and trophic levels. In contrast, the *JEDI* marker approach enables a unified perspective by simultaneously targeting all cellular life in a single PCR reaction. As demonstrated in our application examples, this can provide unique insights, such as the proportional shifts in domain-level diversity within and across ecosystems, that would be unattainable with traditional approaches. By capturing such patterns, the *JEDI* marker approach not only offers a powerful means to advance our understanding of the extent and structure of biodiversity, but also the associations and dynamics that underpin ecological processes and shape ecosystem function and resilience.

Although the *JEDI* marker approach can serve to overcome many of the challenges and limitations in existing biodiversity assessments, it is not without caveats. As presented in two of the example applications, the capacity to detect low-abundance populations, which can disproportionately contribute to ecological processes^70^, is dependent on sequencing depth. As prokaryotic organisms dominate the diversity of most environments, this can have a particularly pronounced effect on the detection of eukaryotic diversity. However, we demonstrate that increasing the sequencing depth improves the extent of eukaryotic diversity recovered (Fig. 5c, Supp. Fig. 4), as well as that of less abundant prokaryotes, as evidenced by the flattening of ASV rarefaction curves across seawater, sediment and soil samples (Fig. 4). Thus, achieving a more complete assessment of biodiversity using the JEDI marker may require greater sequencing depths than typically used for taxon -specific primers. Further improvements for detecting larger eukaryotes and their associated microbiomes can also be made at the sampling level, such as integrating size fractionation alongside bulk sampling as typically performed in plankton ecology^71^. Additionally, while we show that the *JEDI* marker approach proved effective in capturing diversity across domains of life in both a unicellular and multicellular holobiont, these examples do not substantiate the generalisability of its use across all host-associated microbiomes. As the amount of host-derived amplicons can vary widely across holobiont systems and complicate the detection of associated microorganisms, protocol refinements—such as selective lysis, host-blocking strategies, or targeted fractionation—may be required to optimise its application under different contexts. Ultimately, the *JEDI* marker approach places microorganisms at the centre of the biodiversity landscape– aligning with the distribution of diversity on Earth–and, although it represents a significant advancement upon the current *status quo* of biodiversity assessments, it will benefit from further optimisations moving forward.

Looking ahead, there are several opportunities to further enhance the quantitativeness and standardisation of the *JEDI* marker approach through methodological and technological developments. Firstly, while we demonstrate that it provides stable, reproducible and comparable biodiversity estimates across biological systems, achieving a universal framework for global biodiversity assessments will require concerted efforts to standardise sampling, DNA extraction and sequencing protocols. Encouragingly, such standardised practices have already been implemented in large-scale sampling initiatives like *Tara* Oceans^72^ and the Earth Microbiome Project^73^, which can serve as valuable blueprints for building future global DNA sequencing-based biodiversity assessments. Second, as it stands, the *JEDI* marker approach enables comparative estimates of biodiversity across samples and environments, but it does not provide accurate quantification of cells or organisms. Although the use of DNA spike-ins has recently been shown to correct for amplification biases^74^ and provide accurate abundances of organisms containing single copies of the SSU rRNA gene^75^, further developments will be required to calibrate quantification across taxa with different rRNA gene copy numbers, which are known to vary in genomes of different taxa^76^ and correlate to cell size or volume^71^. DNA spike-ins also provide invaluable information on other inherent biases in paired-end sequencing workflows, such as the bias against longer DNA fragments that are typically attributed to eukaryotic organisms–a more detailed insight on this and how to correct for it has been recently covered^77,78^. Supporting the potential to improve these quantitative aspects, ongoing advances in sequencing technologies increasingly allow for the recovery of full-length *JEDI* marker regions and complete SSU rRNA gene operons at high-throughput from environmental samples^79,80^. These advances will allow for discriminating more recent evolutionary divergences and further refine biodiversity assessments. Importantly though, data generated using the *JEDI* marker will remain fully compatible with these future methodological and technological advances, ensuring its continued relevance.

In summary, the *JEDI* marker approach provides a unique mix of practicality, objectiveness and comprehensiveness for assessing biodiversity across the tree of life, from local to planetary scales, advancing beyond the species-centric and macroorganism-biased approaches that underpin the current *status quo*. Although we recognise that its effectiveness depends on factors such as sequencing depth, sampling strategies, and additional optimisations for host-associated systems, we argue that it offers an urgently needed tool to rapidly catalogue and track biodiversity in an era of significant global change, laying the foundation for a holistic biodiversity index that can guide conservation strategies and policy decisions.

## METHODS

### In silico comparison of primers

To support a comprehensive *in silico* comparison of diversity captured by different primer pairs (Table 1), we first constructed curated reference datasets. In particular, we focused on creating reference datasets that span the V3-to-V5 region for prokaryotes and V4-to-V9 region for eukaryotes, given that these are the most frequently targeted regions for the respective groups. To achieve this in a primer agnostic way, we leveraged the SILVA SSU REF138.2^33^ alignment. First, we aligned each primer pair to the reference alignment using SINA v1.7.2^81^ and identified the positions of their target sequences. Second, we aligned sequences from the PR2 v.5.0.1^32^ databases to the SILVA SSU REF NR99 database and subsequently extracted all sequences that spanned either the V3-to-V5 or V4-toV9 primer binding regions. This curation step resulted in the formation of the REF-SQ dataset. Following this, we applied a second curation step to retain only those sequences derived from validly named species (i.e. valid binomial nomenclature), leading to the formation of the REF-SP dataset.

For the comparative analysis of coverage, we searched the primer sequences (after generating all possible variants from ambiguous bases) against the REF-SQ and REF-SP datasets using Blast v2.15.0^82^ (parameters: -task blastn-short -word_size 4 -perc_identity 50 -ungapped -dust no -soft_masking false). The resulting hits were filtered to allow for a maximum of two mismatches, but no mismatches in the final three bases, and a requirement for both forward and reverse primer matches to the same reference sequence. The extent of diversity captured was then determined based on the proportion of sequences (REF-SQ) and taxonomic species (REF-SP) covered.

For the assessment of universal primer coverage across eukaryotic organelles, we downloaded complete genomes of mitochondria (n = 18,206) and chloroplasts (n = 13,342) from the NCBI-organelle database and applied the same blast-based analysis as outlined above.

### In silico assessment of JEDI marker variants vs. taxonomic species

To investigate how diversity estimates of *JEDI* marker ASVs compare to those of taxonomic species, we trimmed the respective region from the sequences of the REF-SP dataset and applied a merging and concatenation-based approach. For references with *JEDI* markers <430 bp in length, we extracted the region as a single fragment, as insert sizes below this threshold would be merged through the DADA2^34^ workflow. For references with JEDI markers >430 bp in length, we trimmed the regions into forward and reverse reads of varying lengths, to mimic different lengths that may arise from quality trimming of amplicon paired-end reads. Resulting read lengths ranged from 90 - 220 bp for forward reads and 80 - 210 bp for reverse reads. These reads were subsequently concatenated. Whole and concatenated fragments were combined and subject to clustering at 97% and 100% identity using cd-hit v4.8.1^87^ (parameters: -c 1.0 -M 10000 -g 1 -r 0). The number of clusters was compared to the number of taxonomic species within the REF-SP dataset. In addition, we assessed the distribution of species within and across clusters to determine the potentiality of under- and over-representation of diversity at a tree-of-life- and taxa-specific level.

### Development of the consensus merge pairs approach

To support accurate, robust and reproducible inference of ASVs from JEDI marker amplicon data, we sought to develop and implement a new functionality into an established, automated workflow, nf-core/ampliseq^39^. The nf-core/ampliseq pipeline was developed to ensure FAIR-conformant end-to-end processing and analysis of raw amplicon sequences into ASVs. However, in line with the standard pipelines of DADA2, ASVs are either inferred from merging or concatenating paired-end reads. The standard approach (and the default setting) is to merge reads based on sharing an overlapping region of >12 bp without mismatches, while those that do not overlap are discarded. For JEDI marker amplicon data, this would result in the loss of most 18S rRNA gene sequences. Therefore, we sought to develop a new functionality that leverages both read merging and concatenation with the inbuilt DADA2 function *mergePairs*. The functionality works as follows, i) Align paired-end reads, ii) Merge paired-end reads with an overlap of >12 bp and no mismatches, iii) Concatenate paired-end reads that share an estimated overlap size below the consensus cut off (1/10th percentile of read overlap sizes), iv) Remove remaining reads. Importantly, we also comprehensively evaluated and optimised alignment score thresholds - based on both simulated and real datasets - to prevent the concatenation of sequences that failed merging only due to errors in the overlapping region (see Supplementary Information). The ‘*consensus merge pairs’* approach (Supp. Fig. 6) has now been integrated into the nf-core/ampliseq pipeline (available through the *mergepairs_strategy* parameter; *consensus* option), available in version 2.13 (see Code Availability).

### Sampling and sequencing for “Application example 1: Tracking cross-domain diversity and dynamics within single-cell and multi-cellular holobionts”

For the investigation of an unicellular holobiont, we isolated unicellular phytoplankton of *Rhizosolenia* sp. through an image-based cell sorting method. Specifically, surface seawater samples were collected in March 2023 at the Astan time series (as part of the TREC expedition) using a 20 µm net tow, and filtered over a 200 µm sieve to retain organisms between 20-200 µm. Using the COPAS Vision 500 (UnionBiometrica) available on the Advanced Mobile Laboratory, we dispensed single cells of *Rhizosolenia* into a 96 well plate containing lysis buffer and processed them for metabarcoding using the universal primers. The amplification of the JEDI marker (including heterogeneity spacers) and attachment of Illumina indices was performed through a two-step amplification procedure. The first PCR included 2 µL from each well with 8 µL of master mix and the Pfu-Sso7d polymerase^88^ and was performed on a thermocycler with the following conditions: 2 min denaturation step followed by 35 cycles of 98 °C for 10 s, 65 °C for 20 s, 72 °C for 20 s, and a final extension of 72 °C for 2 min. The second PCR amplification was subsequently carried out to add Illumina indices, which involved using 2.5 µL of product from the first PCR amplification, 6.5 µL master mix and 1µl of barcodes in a thermocycler with the following conditions: 3 min denaturation step followed by 20 cycles of 98 °C for 30 s, 65 °C for 20 s, 72 °C for 30 s, and a final extension of 72 °C for 3 min. PCR products from the second amplification were pooled in equimolar concentrations and sequenced using an Illumina MiSeq platform, generating 2 x 250 bp with 25,979±10,236 paired-end reads per sample.

To investigate a multicellular holobiont system, we sampled the brown seaweed *Ascophyllum nodosum* at monthly intervals between November 2021 to November 2022 at Pleubian (48.849411, -3.078291, Brittany, France). To access the associated epi- and endophytic microbial community, three thallus parts – apex, middle and base – were sampled from four arbitrarily selected individuals. In brief, after freeze-dried and ground each tissue type sample, the DNA extraction was based on CTAB extraction buffer, followed by two different purification steps: NucleoSpin Plant II kit (MachereyNagel, Germany) and AMPure XP Beads (Beckman Coulter, Brea, CA, USA)^89^. We pooled DNA from the three thallus parts from one individual seaweed. PCRs were performed in duplicate using the *JEDI* primers with Q5® High-Fidelity 2X Master Mix (New England BioLabs, MA, USA) and 3.1 ng of DNA as template. PCR cycling conditions were: 30 s denaturation step followed by 30 cycles of 98 °C for 10 s, 52 °C for 30 s, 72 °C for 30 s, and a final extension of 72 °C for 5 min. PCR products were pooled in equimolar concentrations and sequenced using a Illumina NovaSeq 6000 platform, generating 2 x 250 bp paired-end reads. Eight extraction and PCR blanks were included and served as the basis for sequence decontamination in R^90^ using the microDecon package^91^.

### Sampling and sequencing for “Application example 2: Tracking multi-year, cross-domain diversity dynamics in an ocean ecosystem”

Sampling of surface seawater was conducted bimonthly at the SOMLIT-Astan station in the Western English Channel between January 2009-2012 as described previously^55^. In brief, at each sampling time point, 5 L of water was collected using a Niskin bottle and sequentially filtered through a 3 µm polycarbonate membrane filter and a 0.2 µm sterivex filter (Millipore Sterivex). Filters were preserved in 1.5 ml of lysis buffer (sucrose 256 g/L, Tris 50 mM pH 8, EDTA 40 mM) and stored at −80°C until processing. Filters from both size fractions were incubated for 45 min at 37 °C with 100 µl lysozyme (20 mg ml^-1^), followed by 1 h at 56 °C with 20 µl proteinase K (20 mg ml^-1^) and 100 µl SDS 20% before nucleic extraction using a phenol-chloroform method. Nucleic acids from the size fraction > 3 μm were purified using the NucleoSpin PlantII kit (Macherey-Nagel, silica membrane-based purification). For the size fraction 0.2-3 μm, samples were purified and concentrated by ultrafiltration on Millipore Amicon Ultra 4 columns. Purified DNA extracts were subject to two independent PCR amplifications, following a previously outlined protocol^92^, using a Phusion High Fidelity PCR Master Mix and the eukaryotic-targeting V4 primers (TAReuk454FWD1: 5’-CCAGCASCYGCGGTAATTCC and TAReukREV3: 5’ACTTTCGTTCTTGATYRA)^56^ and the universal SSU rRNA gene primers^26^ (Table 1). Although mismatches in the 3’ end of these eukaryotic-targeting V4 primers prevent the capturing of Haptophyta, they have been shown to exhibit a broad coverage of most eukaryotic taxa^29^. Each sample was amplified in triplicates and subsequently pooled, purified and cleaned using a PCR clean-up kit (Macherey-Nagel). Purified amplicons were pooled in equimolar concentrations and sequenced on an Illumina MiSeq platform, generating 2 x 250 bp paired-end reads.

### Sampling and sequencing for “Application example 3: Measuring biodiversity across the land-sea interface”

Samples for this application example were derived from the Traversing European Coastlines (TREC) project. The TREC project encompassed the sampling of soil, sediment and ocean water samples in transects around the European coast. For this study, we selected representative samples from each ecosystem. Soils were extracted through coring (7 cm diameter, 20 cm depth) followed by homogenization and triplicate 30 -40 ml samples were transferred into a DNase/RNase free, 50 ml cryosafe tube and stored in a cold dryshipper. Sediments were extracted through coring (10 cm diameter, 20 cm depth) and triplicate surface samples of 30-40 ml were obtained using a syringe (after overlaying water removal) and placed into a DNase/RNase free, 50 ml cryosafe tube. For shoreline water, samples were collected using a 5 L Niskin bottle deployed just below the surface at a depth of 0.2–0.5 m. The collected water was pre-filtered through 200 µm and 20 µm mesh sieves into carboys before transfer to a mobile laboratory (van). Water samples were then subject to a sequential filtration procedure encompassing 200–20 µm, 20–3 µm and 3–0.2 µm fractions using vacuum filtration. Each fraction was processed in duplicate from 10 L of the initial water sample and immediately placed into a liquid nitrogen dry shipper for preservation. Surface coastal ocean waters were sampled on the *Tara* vessel following the standard sequential filtration protocol of *Tara* Oceans^72^.

DNA from sediment and soil samples was extracted using a protocol combining mechanical lysis (Lysing Matrix E and Vortex), performed in the presence of a nucleic acid preservation buffer (DNA/RNA Shield), with chemical lysis (Genomic Lysis Buffer), followed by an integrated purification step using the Zymo Quick-DNA Fecal/Soil Microbe Midiprep kit. For shoreline and coastal ocean water samples, DNA was extracted using a cryogrinding lysis step followed by purification with Nucleospin columns (Macherey-Nagel, CITE). Extracted DNA was quantified using a Qubit fluorometer.Three independent PCR amplifications were then performed using the Phusion High Fidelity PCR Master Mix with GC buffer (ThermoFisher Scientific), with universal primers targeting the SSU rRNA gene^9^ (Table 1) and eukaryote-specific V9 primers^45^ (Table 1). PCR amplifications were carried out using a thermocycler with the following conditions: an initial denaturation step of 30 seconds, followed by 25 cycles at 98 °C for 10 seconds, 53 °C for 30 seconds, 72 °C for 30 seconds, and a final extension at 72 °C for 10 minutes.The amplified products from the three PCR replicates were pooled and purified using 1x AMPure XP beads (Beckman Coulter Genomics). The quality of purified products was assessed using an Agilent Bioanalyzer with the DNA High Sensitivity LabChip kit and quantified with a Qubit fluorometer. A library was constructed from an equimolar pool of 8 to 12 purified PCR products and sequenced on an Illumina NovaSeq 6000 platform, generating 2 × 250 bp paired-end reads.

### Processing and statistical analysis of JEDI marker data for application examples

The *JEDI* marker amplicon data generated for each of the application examples was processed into Amplicon Sequence Variants (ASVs) using nf-core/ampliseq with the ‘*consensus merge pairs*’ approach. Reads were trimmed before median quality drops below 25 resulting in trimming lengths of 231 and 230 bp of forward and reverse reads respectively for application example 1 (multicellular holobiont) and application example 2 and trimming lengths of 224 and 223 bp of forward and reverse reads respectively for application example 3. To taxonomically classify the generated ASVs, we employed the dada2 *assignTaxonomy* classifier inside nf-core/ampliseq using default settings. Taxonomic classification was performed against both the PR2 v5.0.0 and SILVA v138 databases, with the former being used to label Eukaryota ASVs and the latter for prokaryotic ASVs.

The ASV tables generated for each application example were analysed in R v4.3.2^90^ with all visualisations generated using the *ggplot2* package^93^, unless otherwise stated. The richness of ASVs in each sample was determined as the mean number of ASVs observed during 500 iterations of subsampling (*rrarefy* function of *vegan*^94^) to the lowest read count in each dataset independently. Additional application example-specific statistical analysis performed are as follows:

● Application example 1: ASVs were grouped based on taxonomic classification into host, fungal symbiont, Eukaryota and Bacteria. A single host count was obtained through summing read counts of ASVs assigned to “*Fucus*”, “*Silvetia*”, and “Phaeophyceae”. Counts of ASVs were subsequently normalised by the host ASV count. The dynamics of ASVs assigned to Bacteria and Eukaryota were compared to those assigned to fungal symbionts through Pearson’s correlation analysis using the *cor.test* function, followed by multiple testing correction using the False Discovery Rate method (*p.adjust* function). Only correlations with a coefficient >0.7 or <-0.7 and an adjusted p-value < 0.05 were retained.
● Application example 3: Rarefaction curves were computed through iteratively subsampling the ASV table (*rrarefy* function of *vegan*) and calculating richness across incrementally increasing counts from 0 to 2 x 10^6^ at intervals of 10,000. The shared and unique ASVs observed across ecosystems was determined with the *ComplexUpset* package using a presence/absence matrix as input. For beta-diversity analysis, we computed Bray-Curtis dissimilarities and Canberra distances between samples using the *decostand* function of *vegan,* visualised differences through Non-Metric Multi-Dimensional scaling with the *metaMDS* function of *vegan* and compared the distances generated from the universal and V9 primers using a mantel test (method = “spearman”, permutations = 999).

## CODE AVAILABILITY

Amplicon sequence data generated from the universal primers in this study was processed through the *consensus merge pairs* approach using the nf-core/ampliseq suite with the following forked version, https://github.com/nhenry50/ampliseq/releases/tag/v0.0.3. Scripts to reproduce the analysis presented in this study are available on a public repository: https://gitlab.com/jedi-paper. The ASV tables and other data tables used by the scripts are available on Zenodo: https://doi.org/10.5281/zenodo.16745475. Since the analysis of the amplicon sequence data, the *consensus merge pairs* approach has been directly integrated into the main nf-core/ampliseq workflow, version 2.13. The unicellular holobiont case study was analysed with this version.

## DATA AVAILABILITY

SSU rRNA gene amplicon data generated for this study are deposited at the European Nucleotide Archive under the following project accession numbers: Application example 1 (Algal microbiome) PRJEB8022. Application example 2 (SOMLIT-Astan time-series data): PRJEB89924. Application example 3 (TREC): SAMEA112553566, SAMEA112553567, SAMEA112490973, SAMEA112491003, SAMEA112489552, SAMEA112551184, SAMEA112561539, SAMEA112489504, SAMEA112489502, SAMEA112496619, SAMEA112496616, SAMEA112496615, SAMEA112496721, SAMEA112496724.

## AUTHOR CONTRIBUTIONS

Key: Concept and Design (CD), Generation of data (G), Analysis of data (A), Interpretation of results (I), Code/software development (S), writing the manuscript (W), revising the manuscript (R), Application example writing (AE-W), Application example concept and design (AE-CD)

Taylor Priest (CD, A, I, W, R). Nicolas Henry (G, A, I, S, W, R). Thomas Weber (G, A, S). Laurine Planat (AE-W, G, A, I, R). Coralie Rousseau and Simon Dittami (G, A, I, R). Yi-Chun Yeh (R), David Needham (R), Hans-Joachim Ruscheweyh (R), Fabienne Rigaut-Jalabert (R), Nathalie Simon (R), Sarah Romac (R), Florence Le Gall (R), Thomas Beavis (G, R), Kevin Moog (G, R), Alice Moussy (R), Corinne Da Silva (R), Caroline Belser (R), ECOMAP team (G), TREC expedition team (G), ML team (G), Jan Korbel (R), Raffaele Siano (R), Julie Poulain (G), Patrick Wincker (R), Paola Bertucci (R), Peer Bork (R), Jed Fuhrman (R). Flora Vincent (AE-W, AE-CD, G, I, R). Shinichi Sunagawa and Colomban de Vargas (CD, I, W, R).

## ACKNOWLEDGEMENTS

We thank the captains and crew of the *Neomysis* research ship for their help during sampling at the SOMLIT-Astan station offshore Roscoff. We thank the TREC consortium, the *Tara* Europa consortium and captains and crew of the R/V Tara, the TREC core partners EMBL, the Tara Ocean Foundation, and the European Marine Biological Resource Centre (EMBRC) for their commitment to making the TREC - *Tara* Europa joint expeditions possible. We extend our special thanks to Christian Jeanthon, Amandine Nunes-Jorge, and Damien Guiffant for insightful discussions and support. This research was primarily supported by the BIOcean5D project, funded by the European Union’s Horizon Europe research and innovation program under grant agreement number 101059915. Views and opinions expressed are however those of the author(s) only and do not necessarily reflect those of the European Union. Additional funding came from the NASA grant number 80NSSC25K0407 - Plankton Planet in support of PACE.

## AUTHORSHIP TEAMS

ECOMAP team: Anne Claire Beaudoux, Aurélie Chambouvet, Christophe Six, Benjamin Bailleul, Daniel Vaulot, Fabrice Not, Frédéric Partensky, Juliana Silva Bernardes, Laure Guillou, Laurence Garczarek, Noriko Yamada, Estelle Bigeard, Florence Le Gall, Morgane Ratin, Charlotte Berthelier, Camille Poirier

Mobile Laboratory team: Niko Leisch, Michael Bonadonna, Tina Wiegand, Paulina Cherek TREC expedition team: TBC

## SUPPLEMENTARY INFORMATION

### Development of the consensus merge pairs approach

Our *consensus merge pairs* approach has been designed to assemble a mixture of overlapping and non-overlapping paired-end reads by merging and concatenation, respectively, without pre-partitioning based on taxonomy (Supp. Fig. 6). Merging is a core step in the inference of ASVs from paired-end amplicon reads, typically performed using the function *mergePairs*() from the DADA2 package. By default, this function aligns paired-end reads and merges those that overlap by more than 12 bp without mismatches. Paired-end reads that do not meet these criteria are discarded. Using this approach on *JEDI* marker data would result in the loss of most 18S rRNA gene sequences due to the region extending beyond the length that can be covered by paired-end reads (Fig. 2a and Supp. Fig. 6 b1). To prevent the loss of this information, the *mergePairs*() function provides an option to concatenate paired-end reads that do not overlap (parameter: *justconcatenate*). However, this step should only be applied to paired-end reads that do not overlap, i.e. applying this to 16S rRNA gene sequences would generate artefacts containing duplicated overlapping regions (Supp. Fig. 6 b2).

To overcome these limitations, we developed the *consensus merge pairs* approach, which combines both merging and concatenation in a simple framework (Supp. Fig. 6 b3) to correctly process both overlapping and non-overlapping paired-end reads. This approach works as follows. 1) Align paired-end reads. 2) Merge paired-end reads with an overlap longer than 12 bp with no mismatches (*justconcatenate* = FALSE). 3) Calculate the overlap **percentile cutoff**: the 1/10 th percentile of the overlap sizes of merged paired-end reads. The percentile cut-off is used to account for paired-end reads that may overlap over a few nucleotides by chance. 4) Concatenate the remaining paired-end reads with an estimated overlap size below the percentile cutoff (*justconcatenate* = TRUE). 5) Discard remaining reads (i.e. paired-end reads of 16S rRNA gene sequences with an overlap greater than the percentile cut -off but containing mismatches - detailed in the next section).

### Benchmarking the consensus merge pairs approach

Mismatches and insertions/deletions (indels) in the overlapping region play an important role in whether reads should be merged or concatenated. Therefore, we set out to empirically assess appropriate mismatch and indel weights for the *consensus merge pairs* approach. For this, we investigated paired-end *JEDI* marker amplicon data from the SOMLIT-Astan time-series (application example 2 of this study). Two pools of DADA2 denoised paired-end reads were generated from the raw data using nf-core/ampliseq version 2.8. For the first pool, forward and reverse reads were trimmed at 200 bp (trunclenf = 200 and trunclenr = 200), while for the second pool, reads were trimmed at 220 bp (trunclenf = 220 and trunclenr = 220). Once denoised, forward and reverse reads were compared to SILVA NR 99 version 138.1 using the VSEARCH’s usearch_global command (https://gitlab.com/metabarcoding_utils/taxonomic-assignment) to identify their best hits (most similar reference sequences). Only best hits more than 90% similar were considered. For the sake of clarity, we refer to a pair of reads as eukaryote when the best hits of both reads were assigned to 18S rRNA gene reference sequences >428 bp in length, and we refer to a pair of reads as prokaryote (including organelles, *e.g.* chloroplasts) when the best hits of both reads best hits were 16S rRNA gene reference sequences <=388 bp in length. That way, eukaryote paired-end reads should not overlap and prokaryote paired-end reads should overlap. At this stage, the 200 bp pool was composed of 16,359 and 2,956 paired-end read unique combinations representing 1,554,326 and 167,528 reads for prokaryotes and eukaryotes, respectively. The 220 bp pool was composed of 15,018 and 2,848 paired-end read unique combinations representing 1,509,287 and 162,477 reads for prokaryotes and eukaryotes, respectively.

The two pools of paired-end reads (200 and 220 bp long reads) were processed using the *consensus merge pairs* approach described above with 36 different weight combinations (match = 1; mismatch = -2, -4, -8, -16, -32, -64; indel = -2, -4, -8, -16, -32, -64). For all weight combinations, almost no eukaryote unique pairs (0.14 and 0.11% for 200 bp and 220 bp pools, respectively) exhibited overlapping regions >12 bp with no mismatch, and thus were merged (justconcatenate = FALSE; Supp. Fig. 6 B3). In contrast, a much larger fraction of prokaryote unique pairs met the merging criteria (52.5 and 41.85 % for 200 bp and 220 bp pools, respectively), representing the majority of prokaryotic reads (95.79 and 94.42% for 200 bp and 220 bp pools, respectively). The corresponding **percentile cutoffs** were 22 and 29 bp for 200 bp and 220 bp pools, respectively. Among the remaining reads pairs, we consider eukaryotic ones as correctly processed if they are concatenated by the *consensus merge pairs* approach (*i.e.* estimated overlap < **percentile cutoff**) and the prokaryotic ones as correctly processed if they are discarded (*i.e.* estimated overlap > **percentile cutoff**).

Alignment weights have a strong influence on the ability of the *consensus merge pairs* approach to properly process prokaryotic read pairs (Supp. Fig. 7). That is, depending on the mismatch and indel weights, the correct processing of prokaryotic read pairs ranges from 0.01-100% of cases, while eukaryotic read pairs are correctly processed in 99.7-100% of cases. In addition, the trimming length employed also plays an important role in the correct processing of read pairs, with proportion of correctly processed prokaryotic read pairs being higher with a trimming length of 220 bp. Based on these results, we determined that an optimal indel weight of -4 and a mismatch weight of -2, which results in the correct processing of more than 99.9% of both prokaryote and eukaryote paired-end reads when the trimming length is 220 bp long. These weights have been set as the default values in the *consensus merge pairs* implementation of nf-core/ampliseq (v2.13; *mergepairs_strategy*=consensus), but they can be adjusted depending on the dataset and/or trimming length.

**Supplementary Figure 1.**
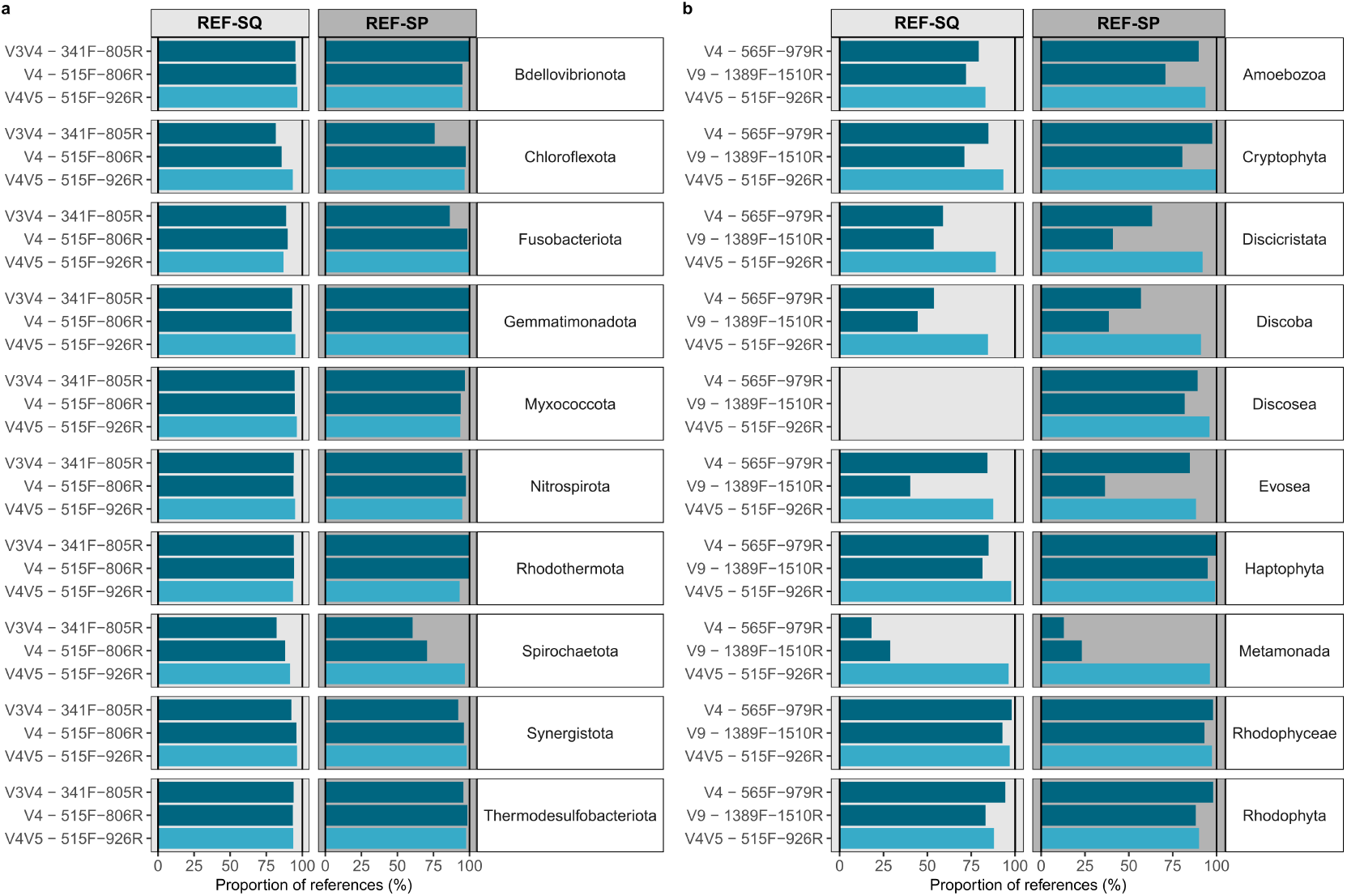
Coverage of primers across taxa within Bacteria and Eukaryota of the REF-SQ and REF-SP dataset. Coverage of primers was determined based on both forward and reverse primer matches to reference sequences (allowing up to two mismatches) within the REF-SQ and REF-SP datasets. The taxa with the largest number of references are presented in Figure 1. This figure illustrates primer coverages across other major (**a**) phyla of Bacteria and (**b**) taxa within Eukaryota.

**Supplementary Figure 2.**
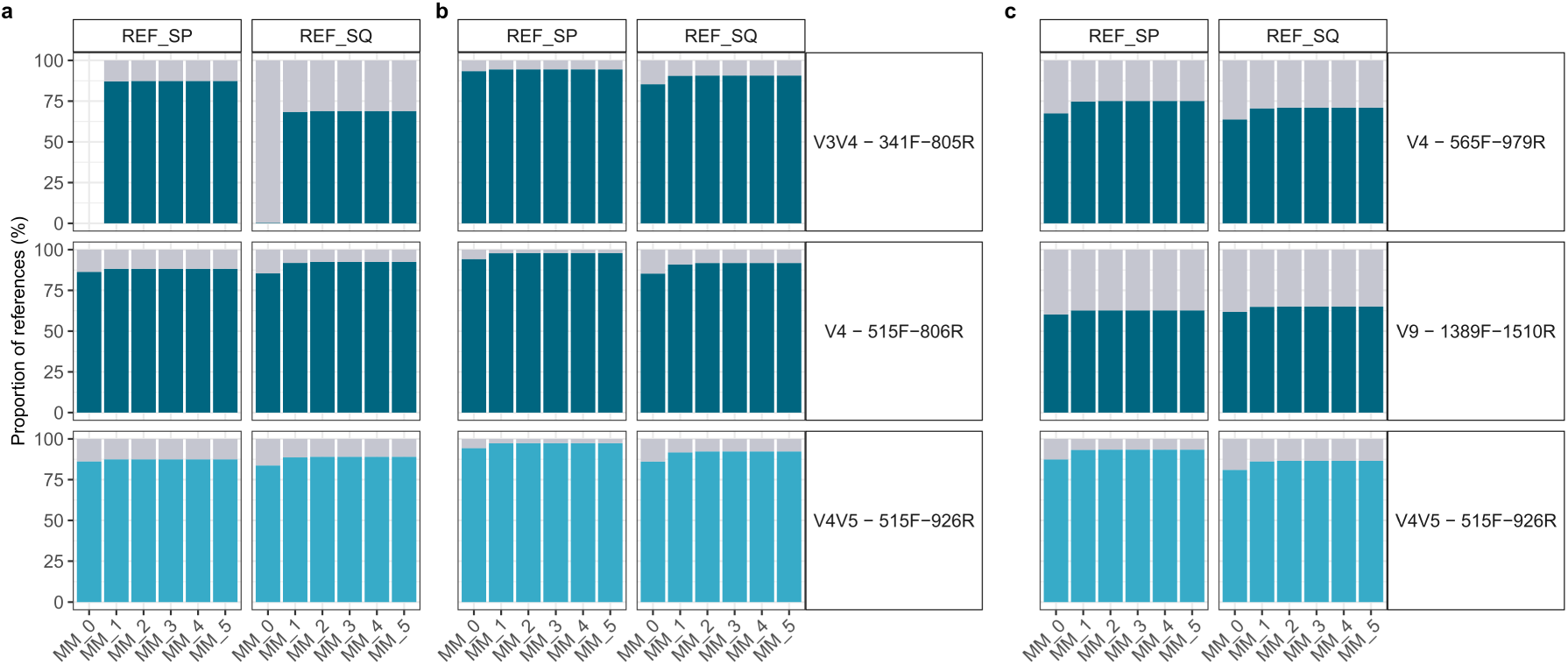
Primer coverage of reference datasets considering different numbers of mismatches. Coverage across **(a)** Archaea, **(b)** Bacteria and **(c)** Eukaryota references in the REF-SP and REF-SQ datasets were assessed considering an increasing number of mismatches in the forward and reverse primers - same number of mismatches in both. However, primer matches to a reference with a mismatch occurring in the last three bases of the 5’ end were excluded, as these may not enable amplification.

**Supplementary Figure 3.**
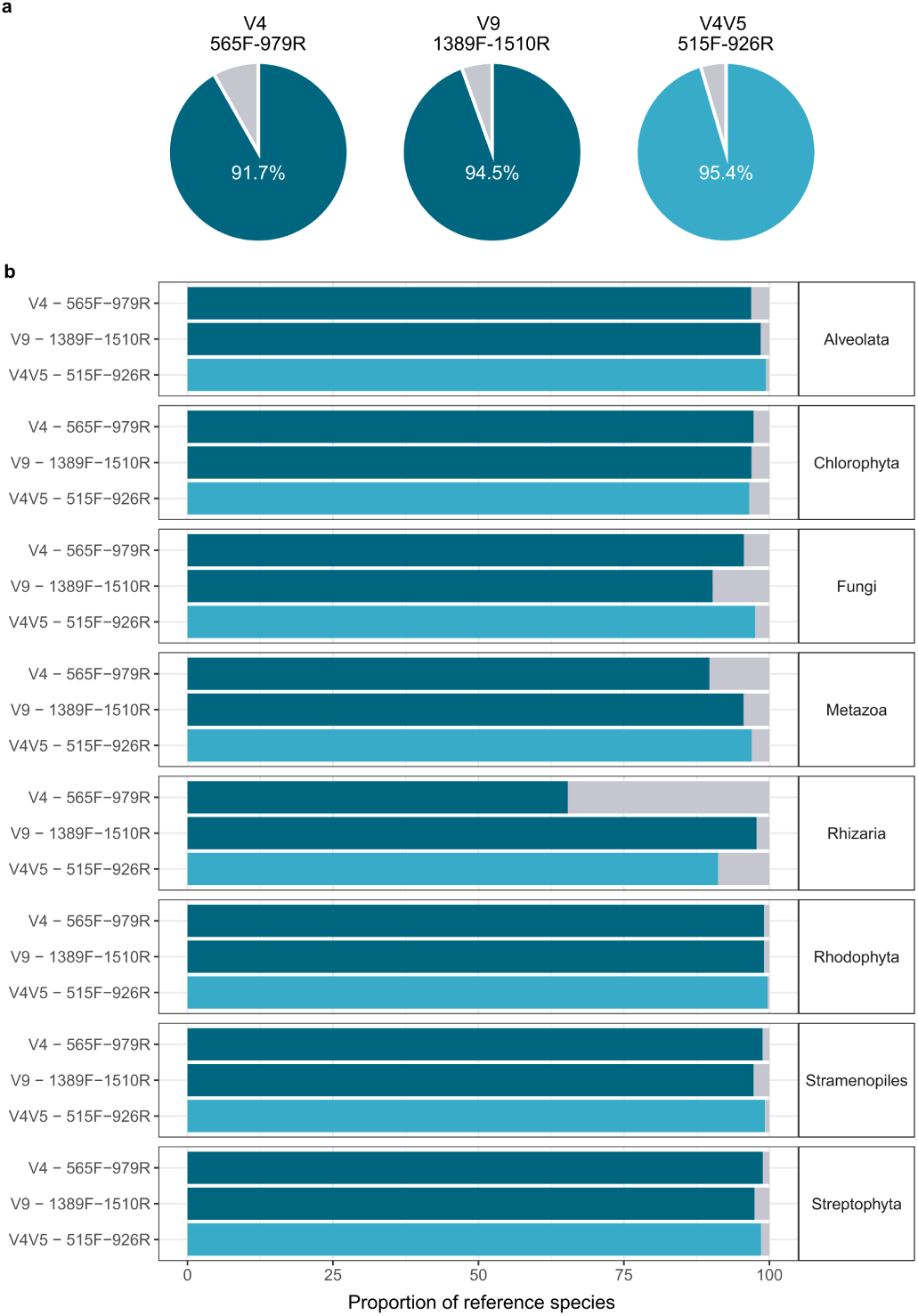
Coverage of forward primers only across major taxa within Eukaryota of the REF-SP. Coverage is based on the proportion of species whose sequences are matched by the forward primer only (allowing up to two mismatches).

**Supplementary Figure 4.**
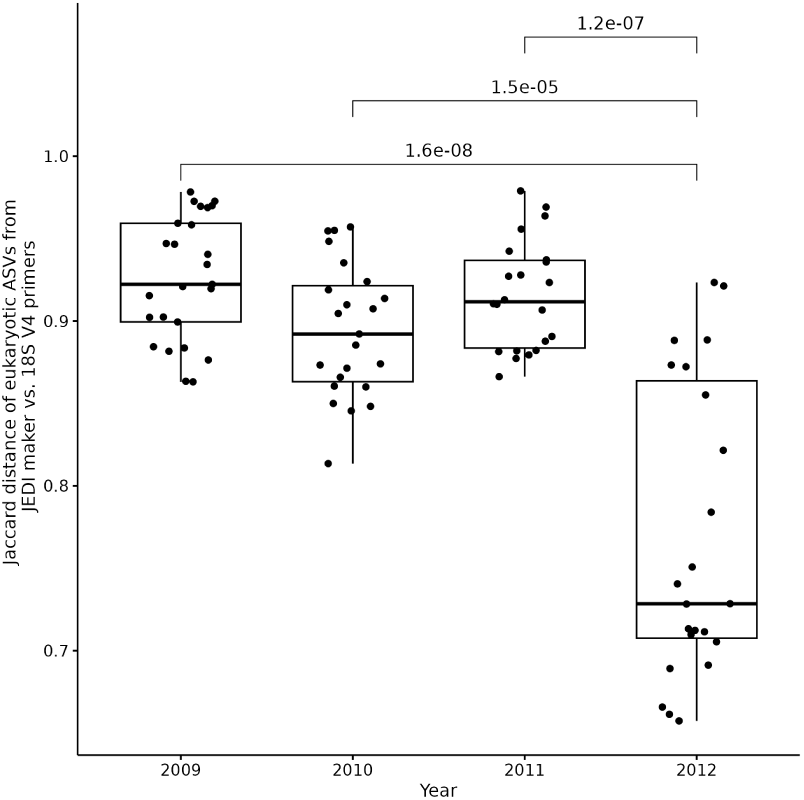
Similarity of the shared sub-ASVs between the eukaryotic ASVs detected using the universal and 18S V4 primers in the Roscoff time-series dataset (for the size fraction >3µm). Eukaryotic ASVs generated from the universal and V4 primers were compared based on their overlapping region - termed sub-ASVs. For each sample, the counts of sub-ASVs were converted to presence/absence and the Jaccard distance calculated between the universal and 18S V4 primers. The boxplots represent the 25th, 50th (median) and 75th percentile of Jaccard distances from samples collected during each annual cycle of the time-series, while the whiskers capture the lowest and maximum Jaccard distances. The Wilcoxon test p-values between 2012 and other years applied on Jaccard dissimilarities are indicated on top of the boxes.

**Supplementary Figure 5.**
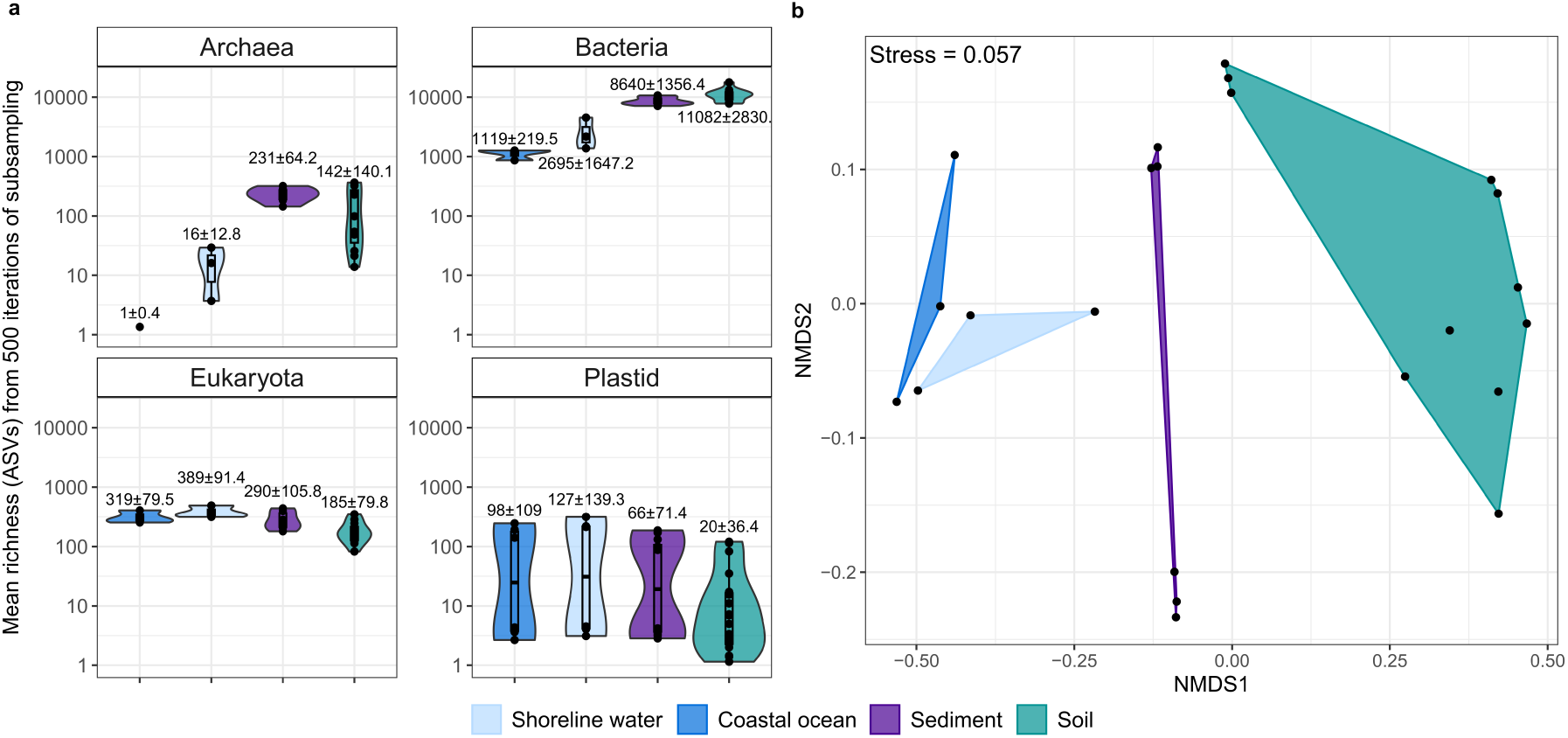
Domain-level richness and composition of communities detected using the *JEDI* marker across different ecosystems. **a)** Mean richness per domain was determined through 500 iterations of subsampling total sample counts to the lowest observed sample count across the dataset. **b)** Non-metric Multi-Dimensional Scaling of Bray-Curtis dissimilarities between sampled communities.

**Supplementary Figure 6.**
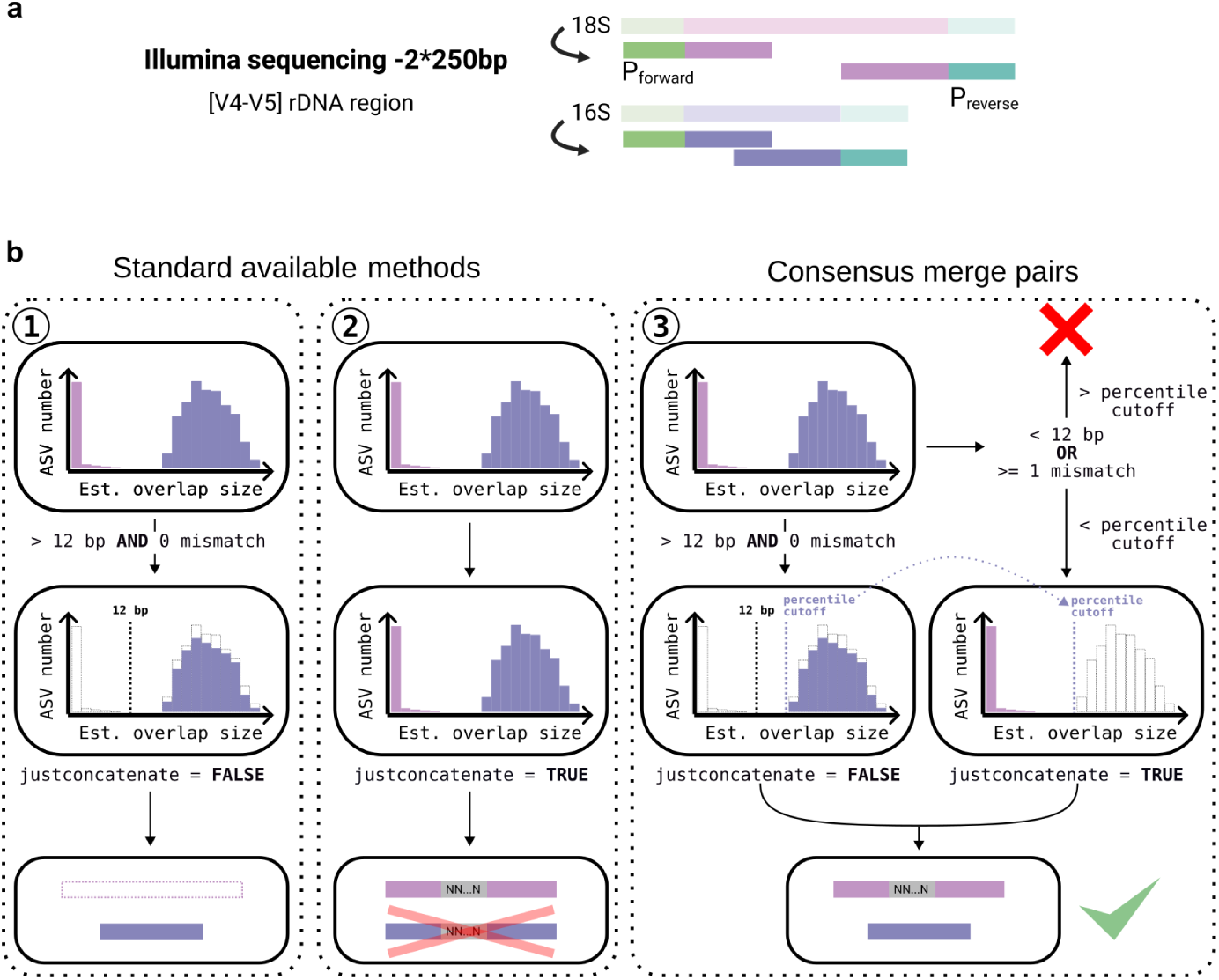
Schematic of the modifications made to a standard amplicon processing pipeline to create the consensus merge pairs approach. Summary of the consensus merge pairs approach (**b3**) in comparison to classical approaches (**b1** and **b2**) applied to a mix of overlapping (16S V4V5) and non overlapping (18 V4V5) paired-end reads of the *JEDI* marker region (**b**).

**Supplementary Figure 7.**
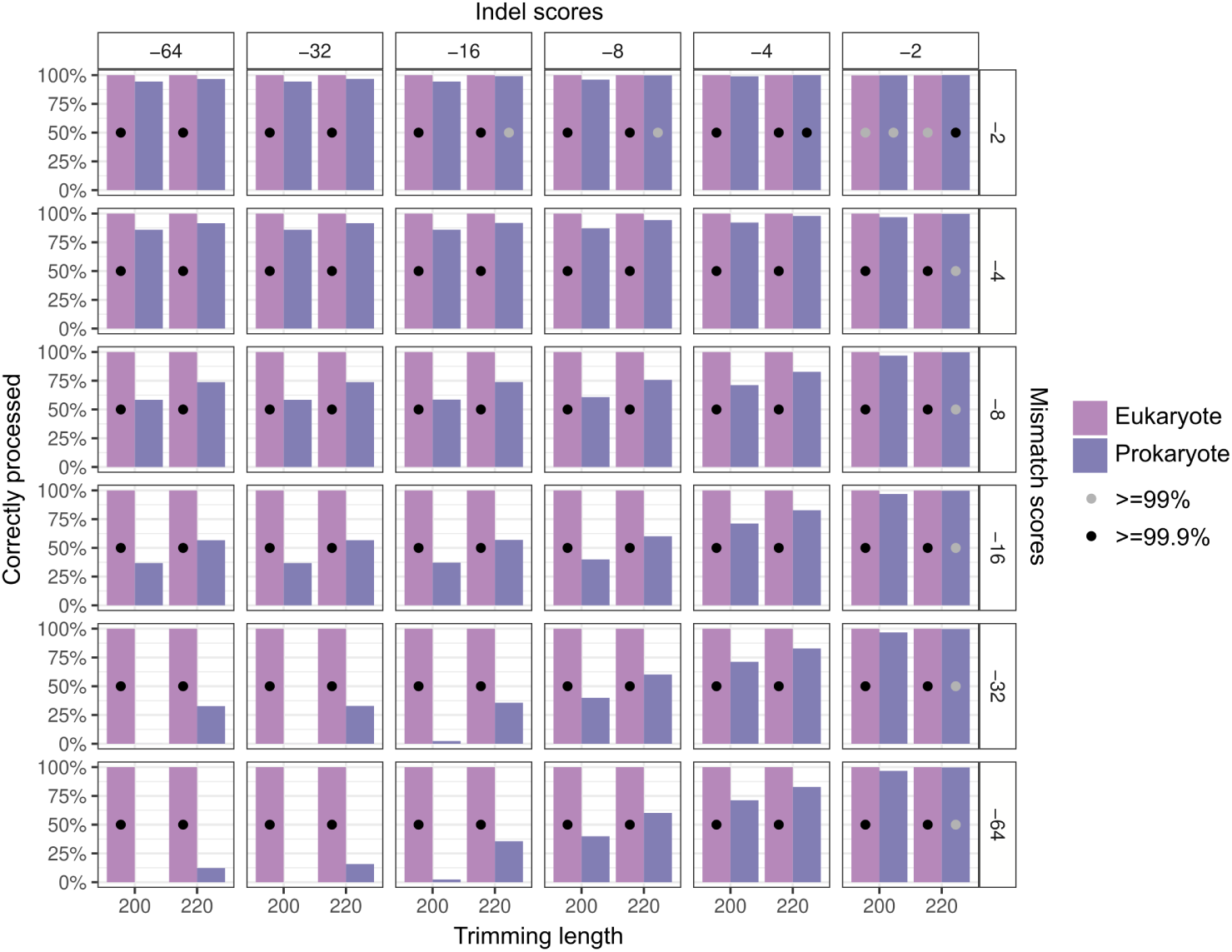
Benchmarking of the mismatch and insertion/deletion weights in the *consensus merge pairs* approach on the SOMLIT-Astan time-series data. The benchmarking of the weights was performed with the same marine time-series data used for application example 2 in this study. The percentage of correctly processed reads (eukaryotes and prokaryotes below and above the overlap percentile cutoff, respectively) is indicated by bars for each read length (200 and 220 bp). Each row corresponds to a mismatch weight and each column corresponds to an indel weight for a total of 36 combinations. Grey and black dots indicate cases when more than 99% (prokaryote) and 99.9% (eukaryote) of paired-end reads were correctly processed.

